# Modulating the cellular context broadly reshapes the mutational landscape of a model enzyme

**DOI:** 10.1101/848010

**Authors:** Samuel Thompson, Yang Zhang, Christine Ingle, Kimberly A. Reynolds, Tanja Kortemme

## Abstract

Protein mutational landscapes are shaped by the cellular environment, but key factors and their quantitative effects are often unknown. Here we show that Lon, a quality control protease naturally absent in common *E. coli* expression strains, drastically reshapes the mutational landscape of the metabolic enzyme dihydrofolate reductase (DHFR). Selection under conditions that resolve highly active mutants reveals that 23.3% of all single point mutations in DHFR are advantageous in the absence of Lon, but advantageous mutations are largely suppressed when Lon is reintroduced. Protein stability measurements demonstrate extensive activity-stability tradeoffs for the advantageous mutants and provide a mechanistic explanation for Lon’s widespread impact. Our findings suggest possibilities for tuning mutational landscapes by modulating the cellular environment, with implications for protein design and combatting antibiotic resistance.

## Introduction

Natural protein sequences are constrained by pressures to maintain required structures and functions within a complex cellular environment, but the key cellular factors are often unknown. In the cellular context, multiple constraints shape the mutational landscapes of proteins, which we define here as the effects on growth of every possible single amino acid mutation in the protein. Deep mutational scanning has been an influential method for determining protein mutational landscapes(Boucher, Bolon, & Tawfik, 2016; Fowler & Fields, 2014) and has provided insights into evolution of new protein functions(McLaughlin, Poelwijk, Raman, Gosal, & Ranganathan, 2012; Stiffler, Hekstra, & Ranganathan, 2015; Wrenbeck, Azouz, & Whitehead, 2017), protein design(Tinberg et al., 2013; Whitehead et al., 2012), functional trade-offs(Klesmith, Bacik, Wrenbeck, Michalczyk, & Whitehead, 2017; Steinberg & Ostermeier, 2016), and adaptation to altered environments(Hietpas, Bank, Jensen, & Bolon, 2013). With a few exceptions(Bandaru et al., 2017; Hietpas et al., 2013; Jiang, Mishra, Hietpas, Zeldovich, & Bolon, 2013; Stiffler et al., 2015), however, these studies find a general tolerance to mutation for residues outside of active sites and binding interfaces(Araya et al., 2012; Boucher et al., 2016; Klesmith et al., 2017; Roscoe, Thayer, Zeldovich, Fushman, & Bolon, 2013; Wrenbeck et al., 2017) that is often explained by the absence of key environmental constraints under the selection conditions(Bandaru et al., 2017; Jiang et al., 2013; Stiffler et al., 2015).

To study the impact of multiple constraints on mutational tolerance during selection, we chose *E. coli* dihydrofolate reductase (DHFR) as a model system. DHFR is an essential enzyme within folate metabolism that reduces dihydrofolate to tetrahydrofolate and is necessary for thymidine production. Using this activity as the basis for an *in vivo* selection assay(Reynolds, McLaughlin, & Ranganathan, 2011), we aimed first to measure a mutational landscape for DHFR and then to determine how a change to the cellular environment might affect the landscape. Because DHFR is known to progress through multiple conformational states during catalysis(Boehr, McElheny, Dyson, & Wright, 2006; Sawaya & Kraut, 1997) (**Figure S1**), we expected the mutational landscape of DHFR to be constrained by the requirement to adopt these different conformations. Moreover, prior work had suggested DHFR is impacted by cellular constraints such as protein quality control (Bershtein, Mu, Serohijos, Zhou, & Shakhnovich, 2013) and the build-up of a toxic metabolic intermediate(Schober et al., 2019). We hence expected deep mutational scanning to reveal a highly constrained mutational landscape for DHFR that would contrast with the mutational tolerance observed in other systems.

## Results

As the basis for our studies, we first sought to establish highly sensitive selection conditions for DHFR function that would be calibrated to DHFR activity and capable of resolving mutants with turnover rates near-to or faster-than wild-type. We anticipated that we would need to control DHFR expression because two prior studies that modified the chromosomal DHFR gene had reported an overall high mutational tolerance under permissive selection conditions that revealed determinants of antibiotic resistance(Garst et al., 2017) and that DHFR expression can be reduced to ∼30% without a growth impact(Bershtein et al., 2013). We used an *E. coli* strain derived from ER2566 with the genes for DHFR and a downstream enzyme, thymidylate synthase, deleted in the genome and complemented on a pACYC-DUET plasmid with a weak ribosome binding site (see **Methods**). To tightly control growth conditions, we performed selections in a turbidostat to maintain the culture in early Log phase growth (**Figure 1A, Figure S2A**). To quantify the effects of DHFR mutations on growth, we calculated selection coefficients from the change in allele frequency over time by deep sequencing of timepoint samples (**Figure 1B**). Under these controlled selection conditions (see **Methods**) we observed a linear relationship between selection coefficient and *in vitro* activity (**Figure 1C**) at cytosolic substrate concentrations(Bennett et al., 2009) for a panel of 14 DHFR mutants (**Table S1**). These results confirm that selection coefficients between −1.5 and 1.0 in our experiment are correlated with DHFR activity over approximately 3 orders of magnitude, and that selection can resolve mutants with higher turnover than wild-type level activity.

**Figure 1.**
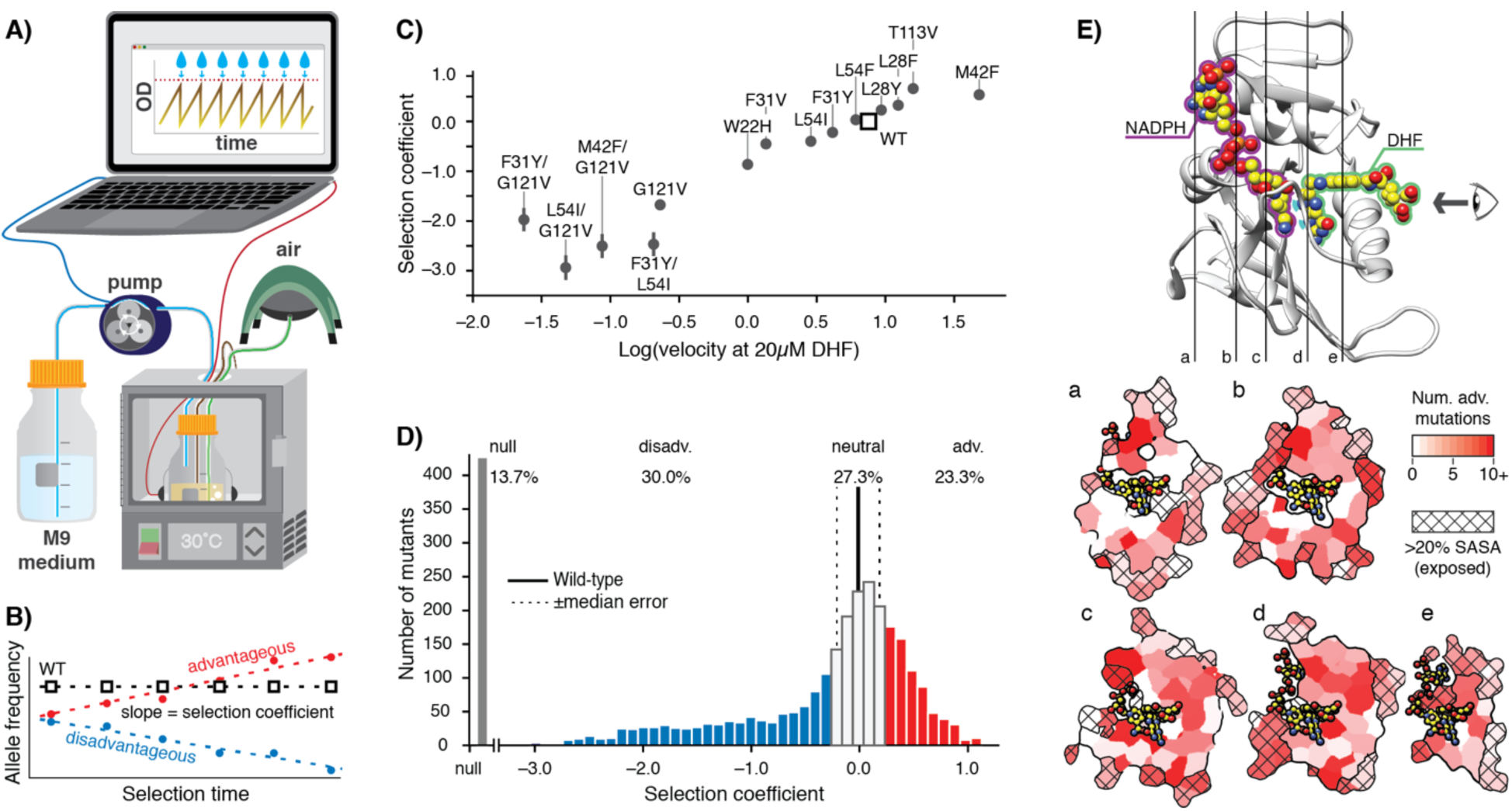
*E. coli* DHFR deep mutational scanning uncovers many advantageous mutations. **A)** Turbidostat schematic. Reoccurring dilutions with fresh medium keep the culture optical density (OD600) below 0.075. **B)** The selection coefficient for each mutant is the slope of the linear regression of allele frequency over time. The wild-type (squares) value is normalized to zero. Advantageous (red) mutations increase and disadvantageous (blue) mutations decrease in frequency. **C)** Selection coefficient as a function of enzymatic activity for purified DHFR point mutants measured *in vitro*. Velocities at 20 µM DHF were calculated from Michalis-Menten parameters. Error bars reflect the standard deviation from 3 biological replicates. **D)** Histogram of selection coefficients. The wild-type value is indicated with a vertical black line. The median standard deviation (**Methods, Figure S2**) is indicated with dashed lines. Mutation are colored as advantageous (red), disadvantageous (blue), neutral (white), or null (grey). **E)** Structural model of DHFR (PDB ID: 3QL3) with cross-section slices (a-e) indicated. The DHF substrate (green) and the NADPH cofactor (purple) are represented by spheres (yellow carbons and heteroatom coloring). An arrow indicates the perspective for each slice. **a-e)** 5 cross-section slices. Color scale indicates numbers of advantageous mutations at each position. Crosshatching indicates residues with >20% solvent accessible surface area.

We next performed deep mutational scanning using the calibrated selection conditions to determine growth effects in biological triplicate for a library of all possible DHFR single point mutants (**Figure 1D, Source data 1**). All pairwise replicates were related with a Pearson correlation R^2^ value of 0.70 and the median standard deviation between replicates for selection coefficients was 0.2 (**Figure S2B-D**). From these data, we define DHFR mutations with selection coefficients of < −0.2 and > 0.2 as disadvantageous and advantageous, respectively. Mutations that were depleted during overnight growth (under less stringent conditions using a supplemented growth medium, see **Methods**) were assigned a null phenotype. As expected, mutations at DHFR positions that are known to be functionally important (M20, W22, D27, L28, F31, T35, M42, L54, R57, T113, G121, D122, and S148) were generally disadvantageous or null mutations (**Figure S3**). These results indicate that our selection assay is a sensitive reporter of functionally important residues and that our results are consistent with previous biochemical characterization of DHFR.

In contrast, the observation of a large fraction of advantageous mutations (red, **Figure 1D**) was unexpected: 737 of 3161 possible variants were advantageous mutations (23.3%), and wild-type DHFR only ranked 1205^th^. In direct measurements of individual growth rates under our selection conditions, the top two DHFR variants (W47L and L24V) led to increases in growth rate of 40 and 76%, respectively, when compared to wild-type DHFR (**Figure S4**). Advantageous mutations were widely distributed over 127 of the 159 positions of DHFR (**Figure 1E**). Furthermore, when we examined the DHFR structure, many of the advantageous mutations appeared to disrupt key side-chain interactions, for example by disrupting atomic packing interactions or surface salt-bridges **(Figure S5)**.

To understand the origins of this counter-intuitive preference for mutations, we looked for cellular factors potentially affecting our mutational landscape. Our selection strain(Anton, Fomenkov, Raleigh, & Berkmen, 2016), like most standard expression strains of *E. coli*, is naturally deficient in Lon protease(Gur & Sauer, 2008) due to an insertion of IS186 in the *lon* promoter region(saiSree, Reddy, & Gowrishankar, 2001). Lon is a major component of protein quality control in *E. coli*(Powers, Powers, & Gierasch, 2012; Sauer & Baker, 2011) responsible for degrading poorly folded proteins. Moreover, Lon had previously been implicated in degrading DHFR unstable variants in *E. coli*(Bershtein et al., 2013; Cho et al., 2015), and deleting Lon in an MG1655 strain of *E. coli* masked the deleterious impact of 2 destabilizing mutations out of a panel of 21 mutants tested in growth experiments at 30°C(Bershtein et al., 2013). Although these 21 mutants were selected for minimal impacts on turnover, we reasoned that the absence of Lon could be responsible for the large fraction of advantageous but potentially destabilizing mutations observed in our selection.

To test this prediction, we reintroduced chromosomal Lon expression under the control of a constitutive promoter in our selection strain, and repeated deep mutational scanning in biological triplicate (**Source data 2**). We refer to the two regimes as +Lon and –Lon selection. The quality of +Lon selection was comparable to that of –Lon selection (**Figure S6**). Consistent with our hypothesis, we observed that the number of advantageous mutations after reintroducing Lon decreased from 737 in –Lon selection to 384 in +Lon selection (**Figure 2A)**, the mean selection coefficient for advantageous mutations decreased from 0.47 to 0.37, and the rank of the wild-type sequence increased by 340 to 865^th^ (**Figure S7**). The median rank of the wild-type residue over all positions decreased from 8 in –Lon selection to 5 in +Lon selection (**Figure S8**).

**Figure 2.**
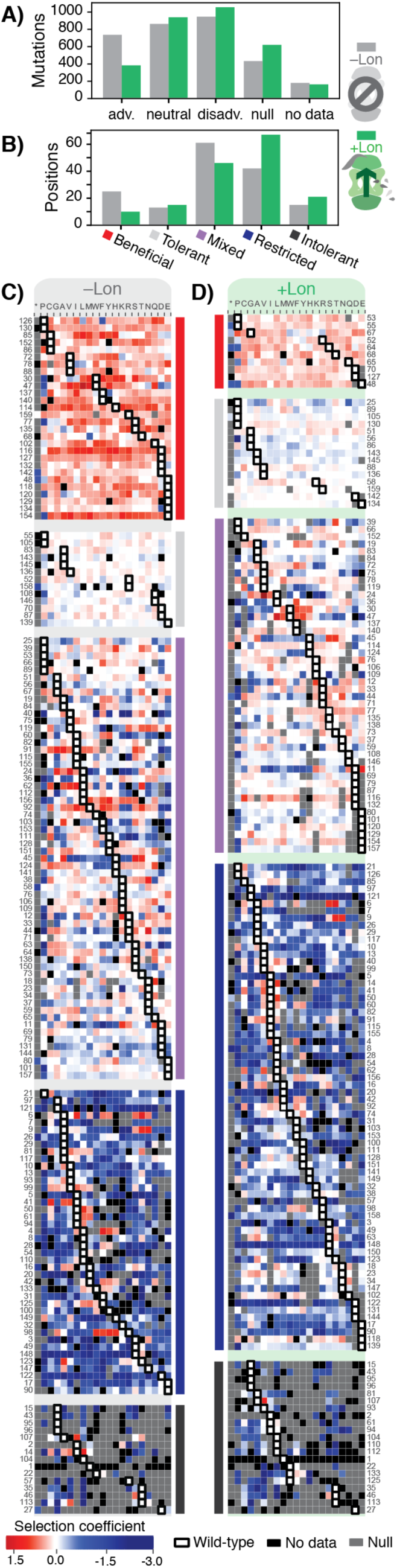
Lon protease expression reshapes the mutational landscape. **A)** Distribution of mutations classified by selection coefficients: 0.2 < =advantageous (adv.), 0.2 > neutral > –0.2, –0.2 => disadvantageous (disadv.), null, and no data (a mutant was not detected in the library after transformation into the selection strain). Grey bars: – Lon selection; green bars: +Lon selection. **B)** Distribution of sequence positions into the 5 mutational response categories: Beneficial, Tolerant, Mixed, Deleterious, Intolerant. Grey bars: –Lon selection; green bars: +Lon selection. **C)** Heatmap of DHFR selection coefficients in the –Lon strain. The heatmap shows advantageous mutations in shades of red, disadvantageous mutations in shades of blue, Null mutations in grey and “No data” as defined in A) in black. Wild-type amino acid residues are outlined in black. Amino acid residues (columns) are organized by physiochemical similarity and indicated by their one-letter amino acid code. An asterisk indicates a stop codon. Positions (rows) are sorted by the wild-type amino acid and grouped by their mutational response category in B) (bars to the right): Beneficial (red), Tolerant (light grey), Mixed (purple), Restricted (blue), or Intolerant (dark grey). **D)** Heatmap for selection in the +Lon strain displayed as in A after reclassifying positions into the mutational response categories (bars to the left) from B).

To examine in more detail how the mutational response of individual residues changes between selection ±Lon, we used a K-means clustering algorithm (see **Methods**) to group all DHFR sequence positions into 5 categories: positions where mutations were generally advantageous (Beneficial), generally neutral (Tolerant), variably advantageous and disadvantageous (Mixed), generally disadvantageous (Restricted), and generally null (Intolerant). Grouping was performed separately for –Lon and +Lon selection (**Table S2**). Comparing the distributions of DHFR positions in –Lon and +Lon conditions illustrates the extensive reshaping of the mutational landscape by Lon (**Figure 2B**). For –Lon selection, 28 positions (17.6%) were classified as Beneficial, where nearly every mutation was preferred over the wild-type residue. In comparison, the number of Beneficial positions decreased to 10 in +Lon selection, with only 3 surface-exposed positions (E48, T68, D127) common between the two Beneficial sets. Simultaneously, the number of Restricted positions increased from 42 to 67 with the reintroduction of Lon into the selection strain (**Figure 2B**). These results support the conclusion that Lon activity broadly penalizes mutations, including a large subset of the advantageous mutations. Overall, the changes upon modulating Lon activity lead to a model in which upregulating Lon increases constraints on DHFR, and the mutational landscape changes from being permissive when Lon is absent to being more restricted when Lon is present (**Figure 2C,D)**.

To analyze the constraints imposed by Lon on the DHFR mutational landscape in structural detail, we defined a Δselection coefficient for each amino acid residue at each position as the difference between the +Lon and –Lon selections (**Figure 3A**). The Δselection coefficient values were most negative at positions in the Beneficial category and at positions with a native VILMWF or Y amino acid residue (**Figure 3B,** excludes Intolerant positions from –Lon selection); overall, mutations at positions with native hydrophobic residues are enriched for negative Δselection coefficients (**Figure S9A**). Strikingly, the mean Δselection coefficient was –0.71 for the 65 buried positions with < 20% side-chain solvent accessible surface area, compared to –0.27 for the 79 exposed positions (**Figure 3C, Figure S9B, Table S2**). These results show that Lon has a broad impact on the mutational landscape throughout the DHFR structure but imposes particularly strong constraints in the DHFR core.

**Figure 3.**
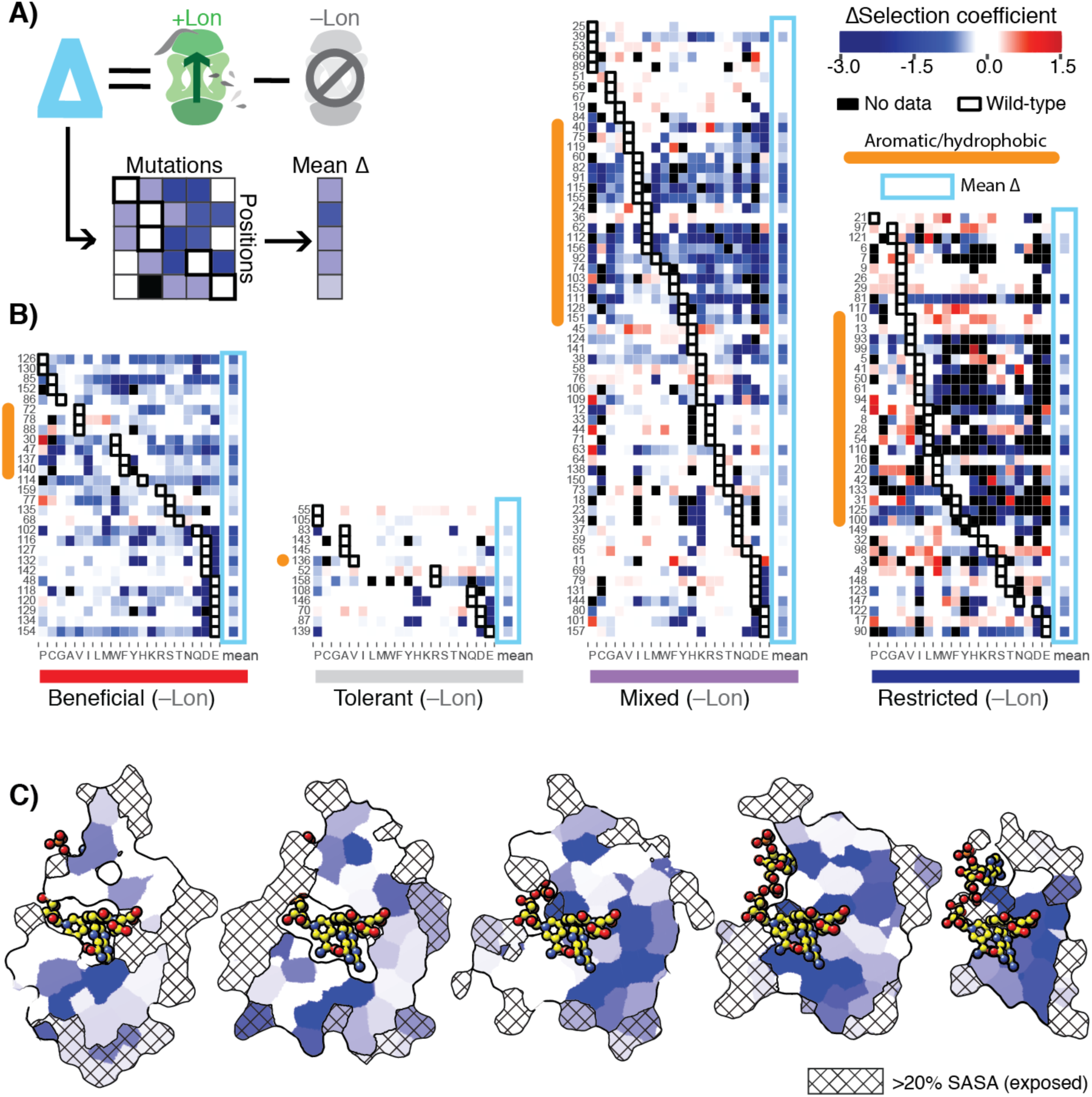
Delta selection coefficients show Lon impact. **A)** Conceptual diagram of Δselection coefficients, calculated as the +Lon selection coefficient minus the –Lon selection coefficient (see **Methods**). **B)** Heatmap of Δselection coefficient values for all positions not classified as Intolerant. Δselection coefficients values between –0.2 and 0.2 are shown in white; Δselection coefficients >0.2 are in shades of red and Δselection coefficients < –0.2 in shades of blue. Amino acid residues (columns) are organized by physiochemical similarity and indicated by their one-letter amino acid code. The mean Δselection coefficient (avg) at each position is shown as a separate column and outlined with a light blue box. Positions (rows) are sorted by the wild-type amino acid and grouped by their mutational response category from the –Lon selection in Figure 2C. Positions with a native VILMWF or Y amino acid are indicated with an orange bar to the left. **C)** Per-position mean Δselection coefficient displayed on the structural model of DHFR. The 5 cross-section slices of the DHFR structure are displayed as in Figure 1E, and the color scale is as in B).

We next sought to more directly test whether advantageous mutations in DHFR destabilize the protein and whether this destabilization could explain their sensitivity to Lon expression. To select specific mutations for *in vitro* tests, we considered all positions with more than one mutation in the top 100 most advantageous mutations. These positions fell into four categories that were each clustered in a hot-spot region in the structure (**Figure 4A,B**, **Figure S10**): 1) exchanges between hydrophobic residues at core positions, 2) disruptions of surface residues on the beta-sheet below the active site, 3) disruptions of polar interactions with the adenine ring of NADPH, or 4) mutations to the active site or M20 loop that controls access to the active site. At these positions, we selected 24 strongly advantageous mutations for *in vitro* characterization. Where possible, we selected two mutations at the same position but with significantly differing Lon sensitivities such that the set had a range of Δselection coefficients from −0.07 to −1.46, with the exception of L24V that had a positive Δselection coefficient. We first confirmed that all selected mutants had *in vitro* activities higher than the wild-type level of activity or were within a two-fold difference (**Figure 4C, Figure S11, Table S3**). We then measured apparent melting temperature (T_m_) values from non-reversible thermal denaturation monitored by circular dichroism spectroscopy, which revealed that many of the advantageous mutations considerably destabilized the protein (**Figure 4D, Figure S12, Table S4**). Moreover and as expected, the Δselection coefficients between +Lon and –Lon selection (**Figure 3**) are correlated with T_m_(**Figure 4D**), except for mutations near the active site. Strikingly, when we compare different mutations at the same position, the change in Δselection coefficients (i.e. Lon sensitivity) correlates with the change in T_m_ values (**Figure 4E**). These results indicate that the selected advantageous mutations are typically destabilizing and that destabilization is correlated with Lon sensitivity. One possible explanation for the selection advantage of destabilizing mutations is that these mutations promote breathing motions that accelerate product release, which is rate limiting for wild-type DHFR at neutral pH(Oyen et al., 2017) and for a hyperactive DHFR mutant with a 7-fold increase in k_cat_(Iwakura et al., 2006).

**Figure 4.**
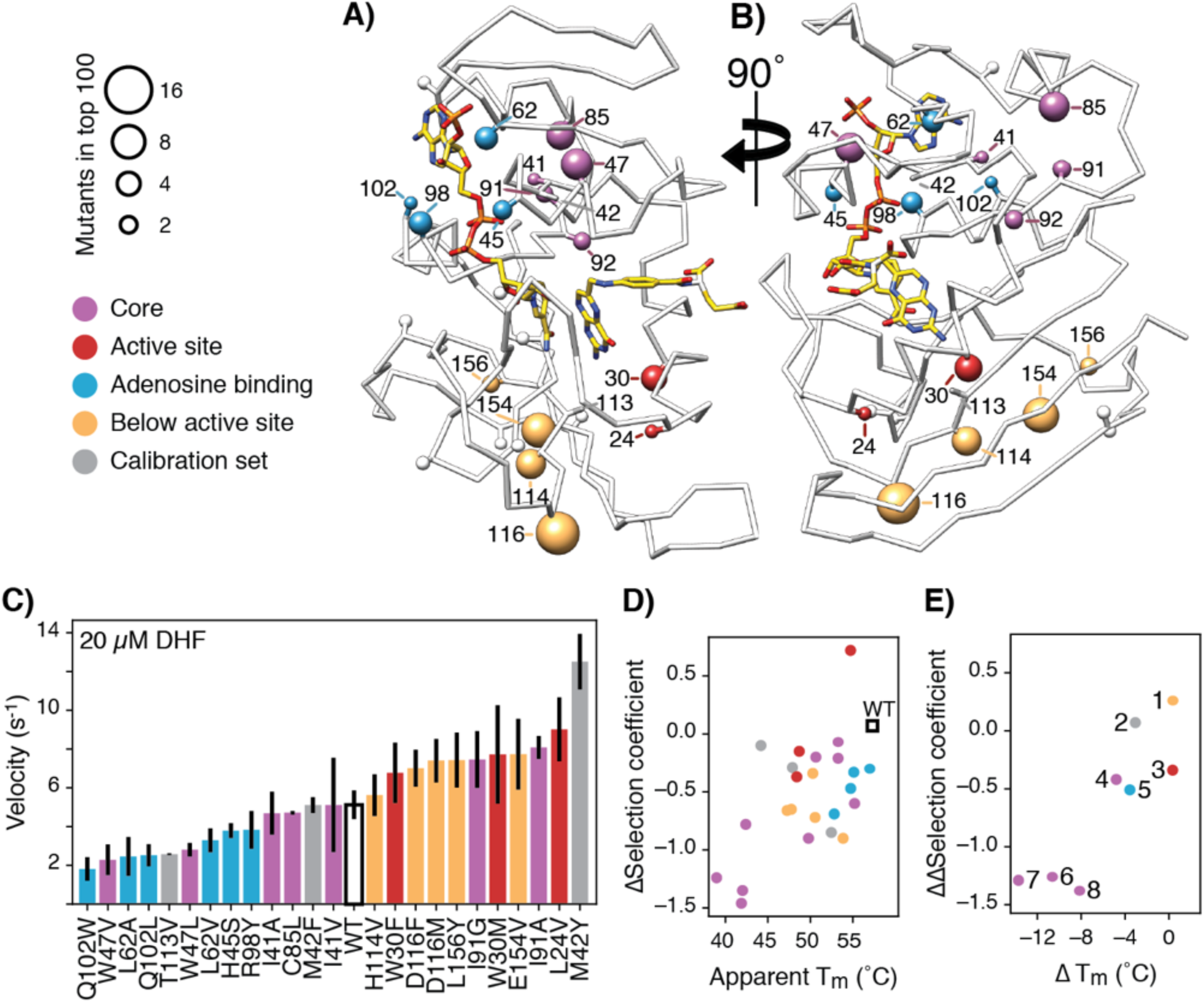
Advantageous mutations are generally destabilizing. **A)** DHFR structure with mutational hot spots. For positions with 2 or more top 100 advantageous mutations, the beta carbon is depicted as a sphere scaled according to the number of top mutations. For mutants selected for *in vitro* characterization, the beta carbon is colored according to its location in the DHFR structure: core (purple), surface beta-sheet (gold), proximal to the adenine ring on NADPH (blue), or proximal to the active site and M20 loop (red). Positions for advantageous mutants from the calibration set are depicted in grey. **B)** The structure from A) rotated 90° clockwise. **C)** *In vitro* velocities of purified DHFR measured at 20 µM DHF. Bars are colored in reference to the hot-spots in A). Error bars represent ±1 standard deviation from three independent experiments (**Methods**). **D)** Correlation between *in vitro* T_m_ values and *in vivo* Δselection coefficients for characterized mutants. Points are colored as in A). **E)** ΔT_m_ values and ΔΔselection coefficient for mutations at the same position. Points representing comparison between mutants are numbered as follows: 1) D116I-M, 2) M42Y-F, 3) W30M-F, 4) I91G-A, 5) Q102W-L, 6) L62A-V, 7) I41A-V, 8) W47V-L.

Taken together, our data indicate that the observed widespread changes in the mutational landscape of DHFR can be explained by a penalty for destabilizing mutations from Lon expression, leading to extensive activity – stability tradeoffs for advantageous mutations. The effect of these two selection pressures is directly observable in the structural arrangement of the mutational response categories (**Figure 5, Figure S13**). In –Lon conditions, mutational responses are arranged in shells around the hydride transfer site(Liu et al., 2013) (**Figure 5A**, top), where the proportion of advantageous mutations increases with increasing distance (**Figure 5B).** This same spatial pattern also holds for +Lon selection (**Figure 5A,** bottom), but it is now superimposed with the additional pressure against destabilizing mutations such that there are no Beneficial positions in the core (**Figure 5C**, **Figure S14**). In contrast, the mutational responses as a function of distance to other DHFR sites (e.g. C5 of the NADPH adenine ring) do not show as strong of a relationship (**Figure S15**). These findings illustrate how the contributions from two constraints – one structural (distance from hydride transfer) and one dependent on cellular context (Lon) – can be dissected from structural patterns in the mutational landscape.

**Figure 5.**
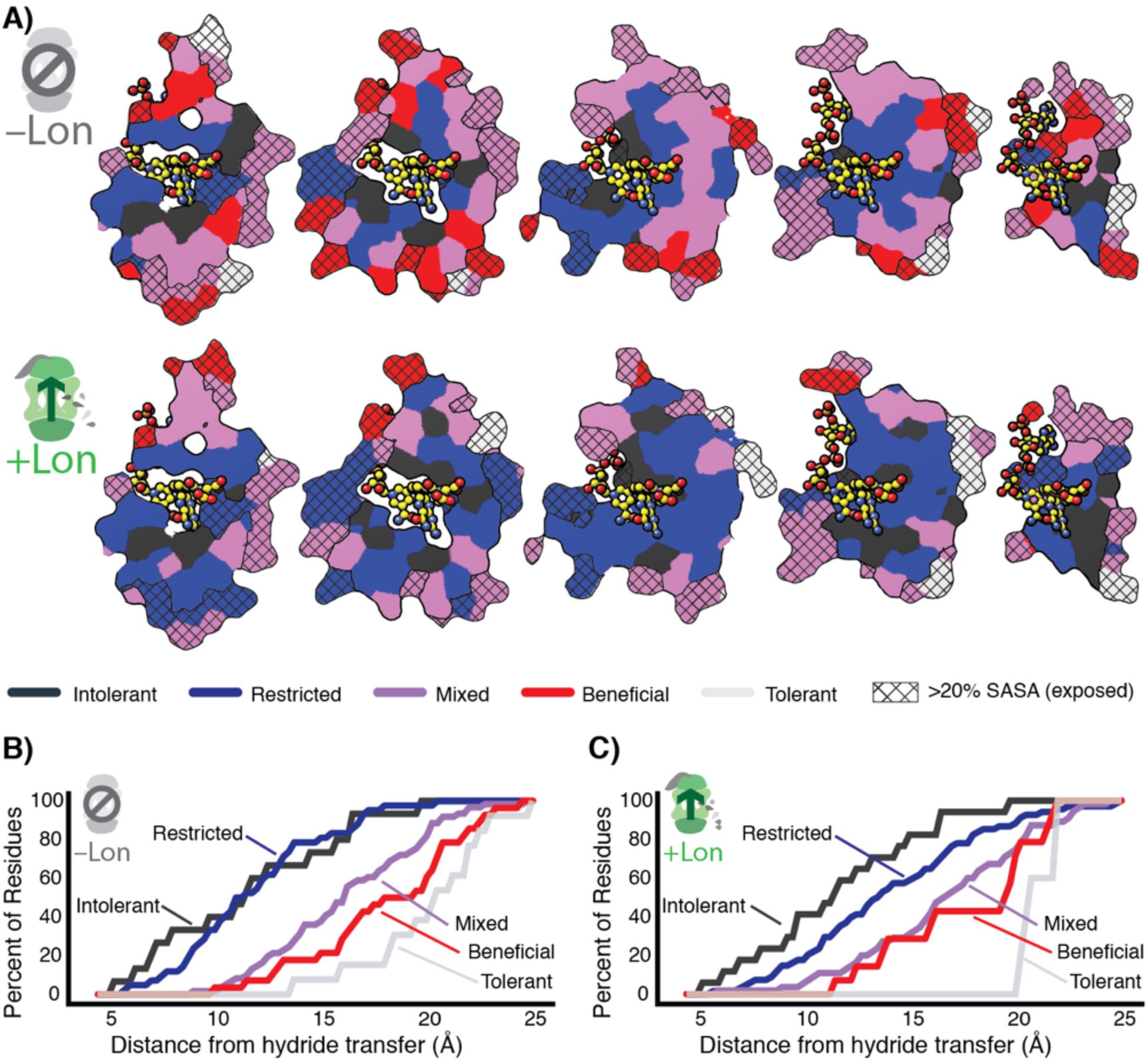
Structural characterization of multiple constraints on the DHFR mutational landscape. **A)** Mutational response categories from –Lon selection (top, categories in Figure 2C) and +Lon selection (bottom, categories as in Figure 2D) colored onto residues and displayed on slices as in Figure 1E). **B)** Relationship between mutational response and distance from hydride transfer for –Lon selection. The percent of positions from each mutational response category are plotted as a function of distance from the site of hydride transfer. Each category colored as in **A**, top). **C)** Relationship between mutational response and distance from hydride transfer for +Lon selection. Each category colored as in **A**, bottom).

## Discussion

The naturally occurring insertion in the Lon promoter in our original selection strain allowed the serendipitous discovery that advantageous mutations are remarkably prevalent throughout the DHFR structure but are also highly sensitive to Lon. The large fraction of advantageous mutations to DHFR appears to conflict with the fixation of the wild-type DHFR sequence during evolution. However, screening DHFR variants under calibrated selection conditions (such as defined temperature, medium, and growth kept in early log phase) for a few generations is not expected not recapitulate the natural selection pressures on *E. coli* DHFR on evolutionary timescales. Our selection is intentionally sensitive to mutation because conditions are calibrated to be linearly related to DHFR activity (**Figure 1C**). In contrast, endogenous DHFR is expected to be buffered from mutational impacts. Reducing DHFR expression in *E. coli* does not have an impact until expression is below 30 % of the endogenous level(Bershtein et al., 2013). Nevertheless, new insights can be drawn from the contrast between mutations that are advantageous in our calibrated selection and the wild-type DHFR sequence. The increase in the number of advantageous mutations in the absence of Lon shows that decreasing environmental constraints can substantially modulate the tolerance to mutation in a deep mutational scanning experiment. From this perspective, tolerance to destabilizing mutations in the absence of Lon explains approximately half of the advantageous mutations and approximately two-thirds of the positions categorized as Beneficial (**Figure 2A, B**). Because all B type *E. coli* strains (e.g. BL21) have the same natural Lon deficiency as our selection strain, our results could have implications for other selection experiments performed in these strains such as the *E.coli* Long-Term Evolution Experiment (Tenaillon et al., 2016), or directed evolution strategies that often lead to mutations at positions distal to the active site.

Beyond experiments in B-type *E.coli*, we expect the fundamental principle of tuning trade-offs to play a role in other experimental systems. Prior work has illuminated the impact of chaperones on the effect of mutations, such as for GroEL in bacteria(Tokuriki & Tawfik, 2009) and for Hsp90 in eukaryotic cells that has been shown to buffer the phenotypic impacts of deleterious mutations(Queitsch, Sangster, & Lindquist, 2002). Our results highlight an opposite key role for the protein quality control machinery to tune *in vivo* mutational responses and lead to a model where protease activities add constraints to the mutational landscape and chaperones relieve them.

The ability to tune multiple constraints could provide a general way of controlling landscapes to drive genes into regions of sequence space that are highly responsive to external pressures. A concrete example of how this principle could be applied is in combinatorial antibiotics. Lon inactivation has been shown to increase resistance to antibiotics(Nicoloff & Andersson, 2013). Switching between compounds capable of inhibiting or activating Lon in combination with DHFR-targeting folate inhibitors such as trimethoprim could serve to variably promote destabilized resistance mutants when Lon is inhibited and then penalize those mutations when Lon is reactivated.

While the power in engineering individual gene sequences is well-recognized, we are only just beginning to explore the potential in engineering the general behavior of local sequence space. We anticipate that further study of tunable constraints will yield a new toolkit for fine control of the landscapes that guide movements through sequence space and enable unexplored engineering applications.

## Materials and Methods

All plasmid and primer sequences are listed in **Source data 3**. Key plasmids were deposited in the Addgene plasmid repository (accession codes are listed in **Source data 3**). All code and python scripts are available at https://github.com/keleayon/2019_DHFR_Lon.git with key input files and example command lines.

### Generation plasmid for in vivo selection assay

The vector bearing DHFR and TYMS for *in vivo* selection (SMT205) was derived from the pACYC-Duet vector described by Reynolds et al (Reynolds et al., 2011). The lac operon upstream of the TYMS gene was replaced with a Tet-inducible promoter. A Tet promoter fragment had been generated with overlap extension PCR and cloned into the pACYC vector (SMT101) at unique AflII/BglII sites to produce SMT201. Selection conditions that resolved increased-fitness mutations were obtained with the SMT205 plasmid where the DHFR “AAGGAG” ribosome binding site (RBS) was replaced with “AATGAG” based on prediction from the RBS calculator(Salis, Mirsky, & Voigt, 2009) using inverse PCR. Briefly, PCR reactions were set up using 2x Q5 mastermix (NEB, cat# M0492), 10 ng of plasmid template, and 500 nM forward and reverse primers. PCR was performed in the following steps: 1) 98°C for 30 seconds, 2) 98°C for 10 seconds, 3) 57-63°C for 30 seconds, 4) 72°C for 2 minutes, 5) return to step 2 for 22 cycles, 6) 72°C for 5 minutes. As needed, the annealing temperature (step 3) was optimized in the range of 57-63°C. 25 µL of PCR reaction was mixed with 1 µL of DpnI (NEB, cat# R0176), 1 µL of T4 PNK (NEB, cat# M0201), 1 µL of T4 ligase (NEB, cat# M0202), and 3.1 µL of T4 ligase buffer (NEB, cat# B0202) at 37°C for 2-4 hours. The reactions were then transformed into chemically competent Top10 cells and plated on LB agar plates with 35 µg/mL chloramphenicol (Fisher BioReagents, BP904, CAS: 56-76-7, 35 mg/mL in ethanol). The plates were incubated overnight at 37°C. Single colonies were picked and used to inoculate 5 mL of LB medium (10g Bacto-tryptone (Fisher BioReagent, cat# BP1415, CAS: 73049-73-7), 5 g Bacto-yeast extract (BD Difco, cat# 212720, CAS: 8013-01-2), 10 g NaCl (Fisher BioReagents, cat# BP358, CAS 7647-14-5), 0.186 g KCl (Sigma, cat# P9541, CAS: 7447-40-7), volume brought to 1 L with MilliQ water, autoclaved) + 35 mg/mL chloramphenicol. Cultures were incubated overnight in 14 mL plastic culture tubes (Falcon, cat# 352059) at 37°C under 225 rpm shaking. Pellets were collected by centrifugation at 3500 rpm for 10 minutes at 4°C in a swinging-bucket centrifuge (Beckman Coulter, Allegra X-12R) and miniprepped (Qiagen, cat# 27104). Constructs were confirmed by Sanger sequencing (Quintara Biosciences) by alignment to the template sequence in ClustalOmega.

### Generation of plasmid libraries

Four sublibraries were generated to cover the entire mutational space of *E.coli* DHFR: positions 1-40 (sublibrary1, SL1), positions 41-80 (sublibrary2, SL2), positions 81-120 (sublibrary3, SL3), and positions 121-159 (sublibrary4, SL4). The single point mutant library was performed by multiple parallel inverse PCR reactions to substitute an NNS degenerate codon at every codon in DHFR. PCR primers (**Source data 3**) were phosphorylated in a 20 µL reaction with 1 µL T4 polynucleotide kinase and 1x T4 ligase buffer. Inverse PCR reactions were performed as described above, followed by PCR clean-up (Qiagen, cat# 28104). The cleaned PCR reactions were incubated for 4 hours with 1µL DpnI, 1 µL of T4 ligase, and 3 µL of T4 ligase buffer. PCR reactions were analyzed by gel electrophoresis using a 1% agarose gel in TAE buffer (20 mM acetic acid (Sigma Aldrich, cat#, 695092), 2 mM EDTA (ACROS Organics, cat# AC118432500, CAS: 60-00-4), 40 mM tris, pH 8.5) with 0.01% v/v GelRed (Biotium, cat# 41003), and the product amount was quantified using gel densitometry in the FIJI image processing software package(Schindelin et al., 2012). Samples were pooled stoichiometrically, cleaned once with a gel extraction kit (Qiagen, cat# 28115), and again with a PCR clean-up kit. The pooled and cleaned ligation products were transformed into *E.coli* Top10 cells by electroporation (BioRad GenePulser Xcell, 1 mm path length cuvette (cat# 165-2089), 1.8 kV, time constant ∼ 5 ms) using ∼5 µL to obtain a minimum of 10^7^ transformants as measured by dilution plating on LB-agar plates with 35 µg/mL chloramphenicol. The transformed cells were rescued in SOB medium (20g Bacto-tryptone, 5g Bacto-yeast extract, 0.584 g NaCl, 0.186 g KCl, 800 mL MilliQ water, pH 7.0, volume brought to 1 L with MilliQ water, autoclaved) without antibiotics for 45 minutes at 37°C before culturing overnight in 10 mL SOB medium with 35 µg/mL chloramphenicol. In the morning, glycerol stocks were made by mixing 500 µL of saturated culture with 500 µL of sterile filtered 50% (v/v) glycerol. 5 mL of the culture was used to miniprep the transformed library with a Qiagen miniprep kit.

### Generation of individual point mutant plasmids

Point mutants in all DHFR-containing plasmids were generated via inverse PCR as described above for the generation of SMT205 except that the appropriate antibiotic was matched with the plasmid (**Source data 3**). Library primer sequences (**Source data 3**) were used except that the “NNS” sequence on the forward primer was replaced with the desired codon.

### Generation of ER2566 ΔfolA ΔthyA –Lon and ER2566 ΔfolA ΔthyA +Lon

The *ER2566 ΔfolA ΔthyA –Lon* strain was generated as previously described(Reynolds et al., 2011) and a gift from Prof. Stephen Benkovic. The ER2566 *ΔfolA ΔthyA +Lon* strain was generated from ER2566 *ΔfolA ΔthyA –Lon* by lambda red recombination using Support Protocol I from Thomason et al. 2014(Thomason, Sawitzke, Li, Costantino, & Court, 2014). The pSim6 plasmid bearing the Lamda red genes linked to a temperature sensitive promoter and the pIB279 plasmid bearing the Kan-SacB positive-negative selection marker(Blomfield, Vaughn, Rest, & Eisenstein, 1991) were gifts from Carol Gross. The Kan-SacB cassette was amplified with 2 rounds of PCR using primers with 5’ homology arms for the region upstream of the Lon gene (**Source data 3**). The insertion fragment containing the Anderson consensus promoter(iGEM, 2006) with homology arms for the region upstream of Lon in the ER2566 genome was amplified from primers using overlap extension PCR.

### Plate reader assay for E. coli growth

Growth rates for the selection strains bearing individual DHFR mutants were measured in 96-well plate growth assays as described for one individual mutant. The SMT205 plasmid was transformed via heat shock into chemically competent *ER2566 ΔfolA ΔthyA ± Lon* cells and plated on an LB-agar plate with 30 µg/mL chloramphenicol plus 50 µg/mL thymidine and incubated overnight at 37°C. On the second day, 2 mL M9 medium (1x M9 salts (BD Difco, cat# 248510), 0.4% glucose w/v (Fisher Chemical, cat# D16, CAS: 50-99-7), 2 mM MgSO4 (Sigma Aldrich, cat# 63138, CAS:10034-99-8)) with supplements for deficient folate metabolism (50 µg/mL thymidine (Sigma Aldrich, cat# T1895, CAS: 50-89-5), 22 µg/mL adenosine (Sigma Aldrich, cat# A9251, CAS: 56-61-7), 1 µg/mL calcium pantothenate (TCI, cat# P0012, CAS: 137-08-6), 38 µg/mL glycine (Fisher BioReagents, cat# BP381, CAS: 56-40-6), and 37.25 µg/mL methionine (Fisher BioReagents, cat# BP388, CAS 63-68-3)) and 30 µg/mL chloramphenicol in a 14 ml culture tube was inoculated with 5-10 colonies scraped from the plate and incubated at 37°C at 225 rpm shaking for 12-14 hours. Biological replicates were obtained from separate inoculations at this step and run on the same plate. All assays were run from fresh transformations. Then, 20 - 50 µL of the previous culture was used to inoculate 5 mL of M9 medium (no supplements) with 30 µg/mL chloramphenicol in a 14 ml culture tube. This fresh culture was incubated for 6 hours at 30°C at 225 rpm shaking. Meanwhile 2 mL of M9 medium with 30 µg/mL chloramphenicol and a transparent 96-well plate were pre-warmed at 30°C. After the 6 hour incubation, the optical density at 600 nm (OD600) of the culture was measured on a Cary 50 spectrophotometer over a path of 1 cm. This early log-phase culture was diluted to an OD600 = 0.005 in the 2 mL aliquot of warmed M9. 200 µL of the dilute culture was pipetted into a well in the 96-well plate. Technical replicates were obtained by dispensing the same dilute culture into multiple wells. Wells were covered with 50 µL of mineral oil (Sigma Aldrich, cat# M5904, CAS: 8042-47-5) using the reverse pipetting technique. The plate was then incubated for 20-48 hours at 30°C in a Victor X3 multimode plate reader (Perkin Elmer). Every 10 minutes, the plate was shaken for 30 seconds with an orbital diameter of 1.8 mm under the “normal” speed setting. Then, the absorbance at 600 nm (ABS600) was measured for each well. Growth rates were calculated from the slope of Log2(ABS600 – ABS600_t=0_) for ΔABS600 in the range of 0.015 – 0.04 using an in-house python script.

### Deep mutational scanning experiments

Competitive growth under selection for DHFR activity was performed in a continuous culture turbidostat (gift of Rama Ranganathan) as described below for a single sublibrary. Sublibraries of DHFR single point mutants were transformed via electroporation as described above into electrocompetent *ER2566 ΔfolA ΔthyA ± Lon* cells using approximately 50 ng of plasmid DNA and 80 µL of competent cells with a transformation efficiency of 10^8^ cfu/ng (based on testing with 10 ng of pACYC plasmid DNA). Immediately after electroporation, the cells were rescued with 2 mL of SOB medium with 50 µg/mL thymidine warmed to 37°C. The rescue culture was incubated at 37°C for 45 minutes at 225 rpm shaking. After the rescue step, 4 µL of the rescue medium (1/500 of the rescue volume) was serially diluted in 10-fold increments. Half the volume of each dilution (1/1000 – 1/10^7^ of the rescue volume) was plated on an LB-agar plate with 30 µg/mL chloramphenicol plus 50 µg/mL thymidine and incubated overnight at 37°C. The colonies were counted the following morning to check for a minimum of 1,000x oversampling of the theoretical diversity in the library (∼10^6^ transformants for each sublibrary). Meanwhile, the larger portion of the rescue medium was mixed with 4 mL of SOB medium with 45 µg/mL chloramphenicol (1.5x) plus 50 µg/mL thymidine warmed to 37°C. This 6 mL culture was incubated for 5-6 hours at 37°C at 225 rpm shaking in a 14 mL culture tube. After incubation, the culture was pelleted by centrifuging for 5 minutes at 3000 rpm at room temperature in a swinging bucket centrifuge. The cells were resuspended in 50 mL of supplemented M9 medium + 30 µg/mL chloramphenicol and incubated for 12-14 hours at 37°C at 225 rpm shaking in a 250 mL flask. In the morning, 150 mL of supplemented M9 medium + 30 µg/mL chloramphenicol in a 1 L flask was inoculated with 15 mL of the overnight culture. This pre-culture was incubated at 30°C for 4 hours at 225 rpm shaking. After 4 hours, the pre-culture was centrifuged at 3000 rpm for 5 min at room temperature in a swinging bucket centrifuge, and the OD600 was measured to ensure that the culture did not grow beyond early-mid log phase (OD600 ∼ 0.3). The supernatant was decanted, and the pellet was resuspended in 30 mL of M9 medium. Pelleting and resuspension were repeated for a total of 3 washes to remove the supplemented medium. After 3 washes, the OD600 was measured for the resuspended pellet using a 10-fold dilution to stay in the linear range of the spectrophotometer.

The washed pellet was then transferred to the growth chamber of the turbidostat (a 250 mL pyrex bottle) containing 150 mL of M9 medium with 50 µg/mL chloramphenicol. Selection experiments were performed with 2 of the 4 sublibraries at a time (two repeats of SL1-SL2 and SL3-SL4, and one repeat of SL1+SL3 and SL2+SL4 for a net of biological triplicates for every codon in the gene), and the resuspended pellet from each library was diluted in the initial culture to an OD600 = 0.035. Mixing and oxygenation was provided by sterile filtered air from an aquarium pump. Every 60 seconds, the aquarium pump was stopped, and the optical density of the culture was read by an infrared emitter-receiver pair. The ADC (analog-to-digital converter) of the voltage over the receiver was calibrated against a spectrophotometer to convert the signal into an approximate OD600. The cells were grown at 30°C with an OD600 threshold of 0.075. When the OD600 of the selection culture exceeded the threshold, the selection culture was diluted to OD600 ∼0.065 with 25 mL of M9 medium with 50 µg/mL chloramphenicol, and the additional culture volume was driven through a waste line by the positive pressure of the aquarium pump. At timepoints of t=0, 2, 4, 6, 8, 12, 16, and 18 hours, 6 mL of the selection culture in 2 mL centrifuge tubes was pelleted at 5000 rpm for 5 minutes at 4°C in a microcentrifuge (Eppendorf, 5242R). The supernatant was removed except for the last ∼200 µL, and the tubes were again pelleted at 5000 rpm for 5 minutes at 4°C in a microcentrifuge, and all the supernatant was carefully removed from the pellet. The pellets were stored at −20°C until sequencing.

### Amplicon generation

Amplicons were generated by two rounds of PCR. The first round of PCR amplifies a portion of the DHFR gene from the pACYC plasmid containing 2-3 sublibraries. For quality control templates were 1 ng/µL plasmid solutions and the amplicons covered SL1-SL2 or SL3-SL4. Round 1 PCR reactions were set up using 1 µL of template, 1% v/v Q5 hotstart polymerase (NEB, cat# M0493), 1x Q5 Reaction Buffer, 1x Q5 High GC Enhancer, 200 µM dNTPs, and 500 nM forward and reverse primers. PCR was performed in the following steps: 1) 98°C for 30 seconds, 2) 98°C for 10 seconds, 3) 57°C for 30 seconds, 4) 72°C for 12 seconds, 5) return to step 2 for 16 cycles, 6) 72°C for 2 minutes.

The Round 2 PCR uses primers that attach the Illumina adapters and the i5 (reverse) and i7 (forward) barcodes for sample identification and demultiplexing. Round 2 PCR reactions were set up and run identically to Round 1 reactions except that the template was 1 µL of Round 1 PCR. Round 2 reactions were analyzed by gel electrophoresis using a 1% TAE-agarose gel in TAE buffer with 0.01% v/v GelRed, and the product amount was quantified using gel densitometry in FIJI. Samples were pooled stoichiometrically and cleaned with a gel extraction kit (Qiagen). Because of the risk of contamination from small primer dimers, gel extraction was performed with very dilute samples. Only 20 µL of sample was loaded onto a 50 mL TAE-agarose gel (OWL EasyCast, B1A) with 8 of the 10 wells combined into a single well. The pooled amplicons were then cleaned again with a PCR clean-up kit (Zymogen, cat# D4013) to allow for small volume elution. The final amplicon concentration was measured with a NanoDrop One UV spectrophotometer and by Picogreen assay (Thermo Scientific, cat# P11496).

### Sequencing of deep mutational scanning experiments

Templates for amplicon PCR were prepared from the frozen pellets. The pellets were resuspended in 20 µL of autoclaved MilliQ water and incubated on ice for 10 minutes. The samples were then centrifuged at 15,000 rpm for 10 minutes at 4°C in a benchtop microcentrifuge. 1 µL of the supernatant was used as template in the amplicon generation protocol for sublibraries described above. The amplicons were sequenced on an Illumina NextSeq using a 300-cycle 500/550 high-output kit. Because of the limitations in the number of sequencing cycles on the Illumina NextSeq, the full amplicon was not sequenced for amplicons containing non-adjacent sublibraries (SL1+SL3, and SL2+SL4). Reads were demultiplexed into their respective selection experiment and timepoint using their TruSeq barcodes. Paired end reads were joined using FLASH(Magoc & Salzberg, 2011). For amplicons with adjacent sublibraries (SL1-SL2 and SL3-SL4), the joined reads were kept. For amplicons with distal sublibraries (SL1+SL3 and SL2+SL4), the unjoined reads were kept. Reads from all lanes of the Illumina chip were concatenated and raw counts of DHFR mutants were obtained from these reads.

Reads on the Illumina NextSeq (two-color chemistry, LED optics) generally have lower quality scores than reads from the Illumina MiSeq (four-color chemistry, laser optics). This lower quality leads to a background signal. This background was estimated from a WT sample. The median + 1 standard deviation value of background count was subtracted from every allele and the alleles were translated into the amino acid sequence, combining synonymous sequences. Counts at each timepoint were only reported for an allele if its frequency was above 2.0 * 10^-5^. Raw counts are reported in **Source data 4-6**.

### Analysis of deep mutational scanning data

Mutant counts were used to generate selection coefficients on our background-subtracted count files with Enrich2 using unweighted linear regression(Rubin et al., 2017). The raw Enrich2 values for each unique selection experiment were combined with a post-processing script. Enrich2 does not calculate selection coefficients for mutants that have no counts at a timepoint, so some selection coefficients were recalculated using only the timepoints before the counts for that allele fell below the cutoff frequency of 2.0 * 10^-5^. Individual selection coefficients were evaluated based on two criteria: noise and number of timepoints. Individual selection coefficients were discarded 1) if the standard error from regression was greater than 0.5 + 0.5 * (selection coefficient) or 2) if there were fewer than 4 timepoints reporting on the mutant. The regression for the fitness value of the mutants from replicate selection experiments to the average values across all experiments was calculated and the fitness values in each replicate were scaled to correct for linear differences in the selection values between replicates. These normalized values were then averaged for the final fitness value. Averaged selection coefficients values were evaluated based on two criteria: the standard deviation of the averaged selection coefficients and the number of replicates. Averaged selection coefficients were discarded 1) if the standard deviation over the normalized replicates was greater than 0.5 + 0.25 * (selection coefficient) or 2) if there were fewer than 2 replicates. In **Source data 1** and **2** the fitness is reported as the mean normalized fitness, the standard error is reported as the combined Enrich2 standard error (from linear regression of timepoints), and the standard deviation is reported as the standard deviation of the biological replicates. The correlation and R-values of normalized replicate experiments and the distribution of standard deviations and standard errors for each mutant is reported in **Figure S1**.

Selection was evaluated by comparing selection coefficients to DHFR velocity from reported Michaelis-Menten kinetics at cytosolic concentrations of DHF (Kwon et al., 2008). Kinetic values are listed in **Table S1**. Based on this calibration, differences between selection coefficients below ∼-2.5 were not considered interpretable, and a floor value of −2.5 was applied to all selection coefficients for the purpose of analysis.

For subtraction to calculate Δselection coefficients, null selection coefficients in +Lon selection were substituted with the lowest measured selection coefficient. Mutations with a null selection coefficient in –Lon selection were assigned a Δselection coefficient of “No data” (colored black). Mutations with “No data” value in either selection condition were also assigned a Δselection coefficient of “No data” here.

### Purification of his_6_-tagged DHFR

DHFR variants were expressed from pHis8 plasmids (KR101/SMT301) for nickel affinity purification as described for one DHFR variant. The plasmid bearing the his-tagged DHFR mutant was transformed via heat shock into chemically competent ER2566 *ΔfolA ΔthyA –Lon* cells, then the cells were plated on LB-agar plates containing 50 µg/mL kanamycin (AMRESCO, cat# 0408, CAS: 25389-94-0, 50 mg/mL in ethanol) and 50 mg/mL thymidine. The plates were incubated overnight at 37°C. The next day 2 mL of LB medium with 50 µg/mL kanamycin was inoculated with a single colony. This culture was incubated overnight at 37°C at 225 rpm shaking. The next day, 25 mL of TB medium (12 g Bacto-tryptone, 24 g Bacto-yeast extract, 0.4% glycerol v/v (Sigma Aldrich, cat# G7893, CAS: 56-81-5), brought to 900 mL with MilliQ water, autoclaved, cooled, mixed with 100 mL sterile filtered buffered phosphate (0.17 M KH_2_PO_4_ (Sigma Aldrich, cat# P0662, CAS: 7778-77-0), 0.72 M K_2_HPO_4_ (Sigma Aldrich, cat# P550, CAS: 16786-57-1))) with 50 µg/mL kanamycin in a 50 mL conical was inoculated with 100 µL of the overnight culture. The culture was grown at 37°C until the OD600 reached 0.5-0.6. Then, the culture was induced with 0.25 mM IPTG (Gold Biotechnology, cat# I2481C100, CAS: 367-93-1, 1M in autoclaved water, sterile filtered) and incubated for 18 hours at 18°C at 225 rpm shaking. The cultures were pelleted by centrifugation at 3000 rpm for 5 minutes at 4°C in a swinging-bucket centrifuge, the supernatant was discarded, and the pellet was resuspended in 4 mL/g-pellet of B-PER (ThermoScientific, cat# 78266) with 1 mM PMSF (Millipore Sigma, cat# 7110, CAS: 329-98-6, 100 mM in ethanol), 10 µg/mL leupeptin (VWR Chemicals, cat# J583, CAS: 26305-03-3, 5 mg/mL in water), and 2 µg/mL pepstatin (VWR Chemicals, cat# J580, CAS: 103476-89-7, 2 mg/mL in water). The lysate was then incubated for 30 minutes with 20 µL of NiNTA resin pre-equilibrated in Nickel Binding Buffer (50 mM tris base (Fisher BioReagents, cat# BP152, CAS: 77-86-1) pH 8.0, 500 mM NaCl, 10 mM imidazole (Fisher Chemical, cat# 03196, CAS: 288-32-4), and then supernatant was removed by pipetting. The resin was washed 3 times for 5 minutes with 1 mL of Nickel Binding Buffer. Then the protein was eluted into 200 µL of Nickel Elution Buffer (100 mM tris pH 8.0, 1 M NaCl, 400 mM imidazole) and dialyzed against DHFR Storage Buffer (50 mM tris pH 8.0, 300 mM NaCl, 1% glycerol v/v) in 3000 Da MW cut-off Slidalyzer dialysis cups (Thermo Scientific, cat# 88401) at 4°C. After 4 changes of dialysis buffer over 24 hours, the protein was aliquoted, flash frozen in liquid nitrogen, and stored at −80°C. Proteins were purified to ∼90-95% purity as judged from PAGE gel analysis.

### In vitro kinetics for DHFR activity

*In vitro* measurements of DHFR activity were carried out by monitoring the change in UV absorbance. For each mutant screened, a purified enzyme aliquot was thawed and centrifuged at 15,000 rpm for 5 minutes at 4°C in a benchtop microcentrifuge. The soluble enzyme was then transferred to a fresh tube, and the concentration was measured by UV absorption on a Nanodrop. Molar concentration of DHFR was calculated using an extinction coefficient of 33585 M^-1^ cm^-1^ at 280 nm for all variants with the following exceptions: 28085 (W30F/M, W47L/M), 35075 (M42Y, R98Y, L165Y), or 39085 (Q102W) M^-1^ cm^-1^. The enzyme was diluted to 555 nM in DHFR storage buffer. A pre-reaction mixture was prepared in MTEN buffer (5 mM MES (Sigma Aldrich, cat# 69889, CAS: 145224-94-8), 25 mM ethanolamine (Sigma Aldrich, cat# E6133, CAS: 2002-24-6), 100 mM NaCl, 25 mM tris base, pH to 7.0) with 55.5 nM enzyme, 111 µM NADPH (Sigma Aldrich, cat# N7505, CAS: 2646-71-1) and 5 mM DTT (GoldBio, cat# DTT25, CAS: 27565-41-9, 1M in water, sterile filtered). The pre-reaction mixture and a micro quartz cuvette (Fisher Scientific, cat# 14-958-103, 10 mm path length, 2 mm window width) were pre-incubated at 30°C. The reaction was started by adding 20 µl of 500 µM DHF (Sigma Aldrich, cat# D7006, CAS: 4033-27-6) in MTEN with 5 mM DTT to 180 µL of pre-reaction mixture. The substrate solution was made fresh from a sealed ampule on the day of the experiment. The reaction was briefly mixed by pipetting and then the reaction was monitored by reading the absorbance at 340 nm with an interval of 0.1 seconds in a Cary 50 spectrophotometer with the Peltier temperature set to 30°C. The reactions were allowed to run to completion to establish the baseline, which was subtracted from the absorbance values. The real-time concentration of DHF was calculated by dividing the normalized absorbance values by the decrease in absorbance at 340 nm for the reaction, 0.0132 µM^-1^cm^-1^, the velocity of the reaction was calculated as the slope of linear regression to a 30 second window with a mean DHF concentration equal to 5, 10, 20, or 30 µM. Final velocities were normalized to enzyme concentration.

Michaelis-Menten kinetics were performed as described above using 1-5 µM DHFR for concentrations of DHFR from 0.5-100 µM. Initial velocities were estimated from linear regression to the absorbance divided by the decrease in absorbance at 340 nm for the reaction, and then they were fit to the Michaelis-Menten equation using the non-linear least squares method in R.

### CD Spectroscopy

Samples for circular dichroism (CD) spectroscopy were prepared at a concentration of 10 µM in a buffer of 150 mM NaCl and 50 mM Tris, pH 8.0. CD spectra acquisition and thermal denaturation was carried out in a Jasco J-715 CD spectrometer using a cuvette with a 2 mm pathlength (Starna Cell Inc., cat# 21-Q-2). For each DHFR variant, a pre-denaturation spectra was recorded between 207 nm and 280 nm where the high tension voltage was below 600 V. Thermal denaturation data was collected at 225 nm with a bandwidth of 2 nm, a response time of 8s, and a resolution of 0.1°C during heating at a rate of 1°C/min. When the curve flattened, the sample was removed from the CD spectrometer and the system was returned to 30°C. The sample was returned to the chamber and allowed to equilibrate for 10 minutes. A post-denaturation spectrum was recorded after equilibration. Between samples, the cuvette was cleaned with sonication in Hellmanex III (Hellma, cat# 2805939) followed by washing with 50% concentrated nitric acid. Thermal denaturation was found to be only partially reversible based on comparisons of spectra recorded before and after denaturation. Thermal denaturation curves were fit to a sigmoidal model for the calculation of an approximate apparent T_m_ for all mutants as previously reported(Smith et al., 2013).

### Structural representation of DHFR

All images of the DHFR structure were prepared with UCSF Chimera, and volumetric representations were prepared using MSMS package(Sanner, Olson, & Spehner, 1996). Solvent accessible surface accessible surface area (SASA) was calculated using the Getarea server(Fraczkiewicz & Braun, 1998) for 4 crystal structures of DHFR (1RX1, 3QL3, 1RX4, and 1RX5) representing different states in DHFR’s catalytic cycle. All models were downloaded from PDB_REDO(Joosten, Long, Murshudov, & Perrakis, 2014). For all positions in DHFR, if the residue had < 20% SASA in any structure, the residue was classified as buried. All other residues were classified as exposed. Burial classification is reported in **Table 2**.

The distance between the positions within each mutational response category and sites within the DHFR structure (hydride transfer site, M20 loop, core of the globular domain, and the beta-sheet surface beneath the active site) were determined using a model of the transition state provided by Phil Hanoian(Liu et al., 2013). The representative atom for the hydride transfer site is the hydride atom in the transition state model. The representative atom for the adenine ring is C5 (C18 in the pdb). The representative atom for the core of the globular domain is the alpha carbon of I41. The representative atom for the beta sheet region is the alpha carbon of D114. For all cases, the distance is defined as the distance between the representative atom and the alpha carbon of the target position.

Mean atom neighbors for each residue on a structure were calculated using an in-house python script. The number of non-hydrogen atoms within an 8 Å shell of each non-hydrogen atom in the structure were counted and averaged for all non-hydrogen atoms at each side chain. These values we calculated for 4 crystal structures of DHFR (PDB IDs: 1RX1, 3QL3, 1RX4, 1RX5) and averaged over the set.

## Acknowledgements

The authors would like to thank Carol Gross, Melanie Silvis, and Byoung Mo Koo for discussion and for providing Lambda red plasmids; Rama Ranganathan and Victor Salinas for supplying parts and expertise for the construction of the turbidostat; Sharon Hammes-Shiffer and Phil Hanoian for providing QM/MM models of the hydride transfer step; Natasha Carli and Jim McGuire at the Gladstone Institute Genomics Core for performing NextSeq 500 sequencing runs with support from the James B. Pendleton Charitable Trust; and Norma Neff, Anna Sellas, and Rene Sit for performing and aiding Miseq and NextSeq 500 sequencing runs at the Chan Zuckerberg Biohub.

## Author contributions

SMT and TK conceived the project idea and experimental approach. SMT, TK, and KAR analyzed experimental data. TK, KAR, and SMT acquired funding. SMT designed and performed the majority of experiments with help from YZ (protein expression and CD spectroscopy) and CI (selection optimization and deep sequencing). CI provided and managed key reagents, strains, and plasmids. TK and KAR provided advice, mentorship, and resources. SMT and TK wrote the manuscript with input from all authors.

## Data and materials availability

All relevant data are in the main text or Supplementary Information and raw sequencing counts are provided in Source Data 4-6. Raw deep sequencing data was deposited to the Sequence Read Archive in entry PRJNA590072. Key plasmids (**Source data 3**) will be available from Addgene upon publication.

