## Supplementary Figures and Tables for "Modulating the cellular context broadly reshapes the mutational landscape of a model enzyme"

**This PDF file includes:**

Figs. S1 to S15
Tables S1 to S4

**Other Supplementary Materials for this manuscript include the following:**

Source data 1 – Selection coefficients for –Lon selection
Source data 2 – Selection coefficients for +Lon selection
Source data 3 – Primers and plasmids
Source data 4 – Raw deep sequencing counts for calibration mutants in –Lon selection
Source data 5 – Raw deep sequencing counts for –Lon selection
Source data 6 – Raw deep sequencing counts +Lon selection

**Figure S1.**

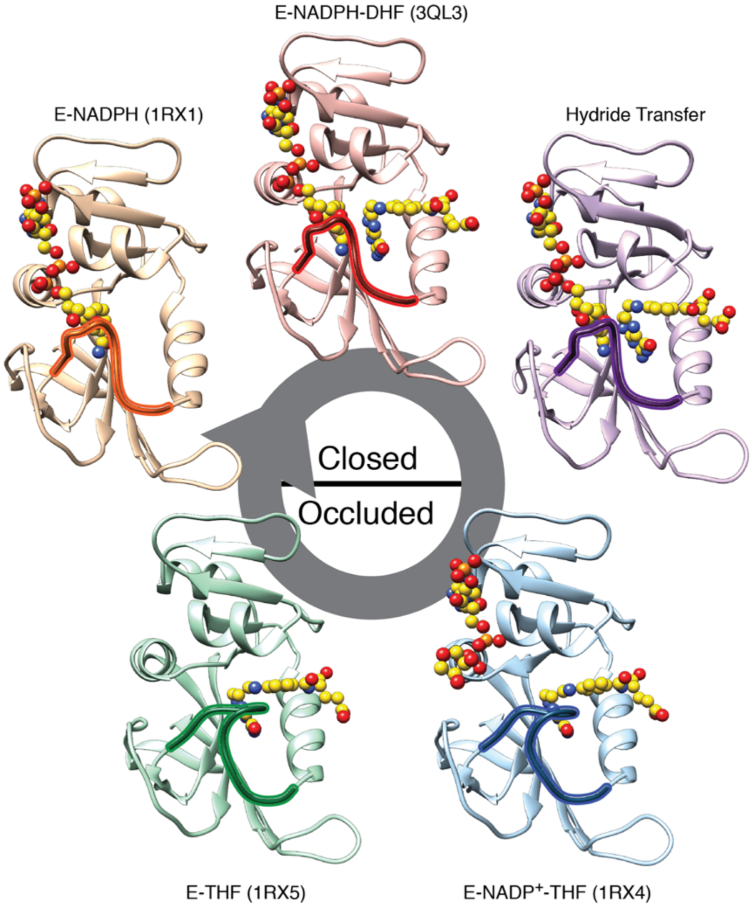


**Fig. S1. Conformations adopted during the DHFR catalytic cycle**. Crystal structures of DHFR (PDB IDs: 1RX1, 3QL3, 1RX4, and 1RX5) and a QMMM model of the hydride transfer step(Liu et al., 2013) represent the conformational states adopted by DHFR over the catalytic cycle. The identity of each state is defined by the identity of the bound ligands (yellow spheres with heteroatom coloring) and the conformation of the M20 loop (outlined) that folds over the active site (closed or occluded). Upper models are in the closed state, and lower models are in the occluded state. All PDBs were downloaded from the PDB_REDO(Joosten, Long, Murshudov, & Perrakis, 2014).

**Figure S2.**

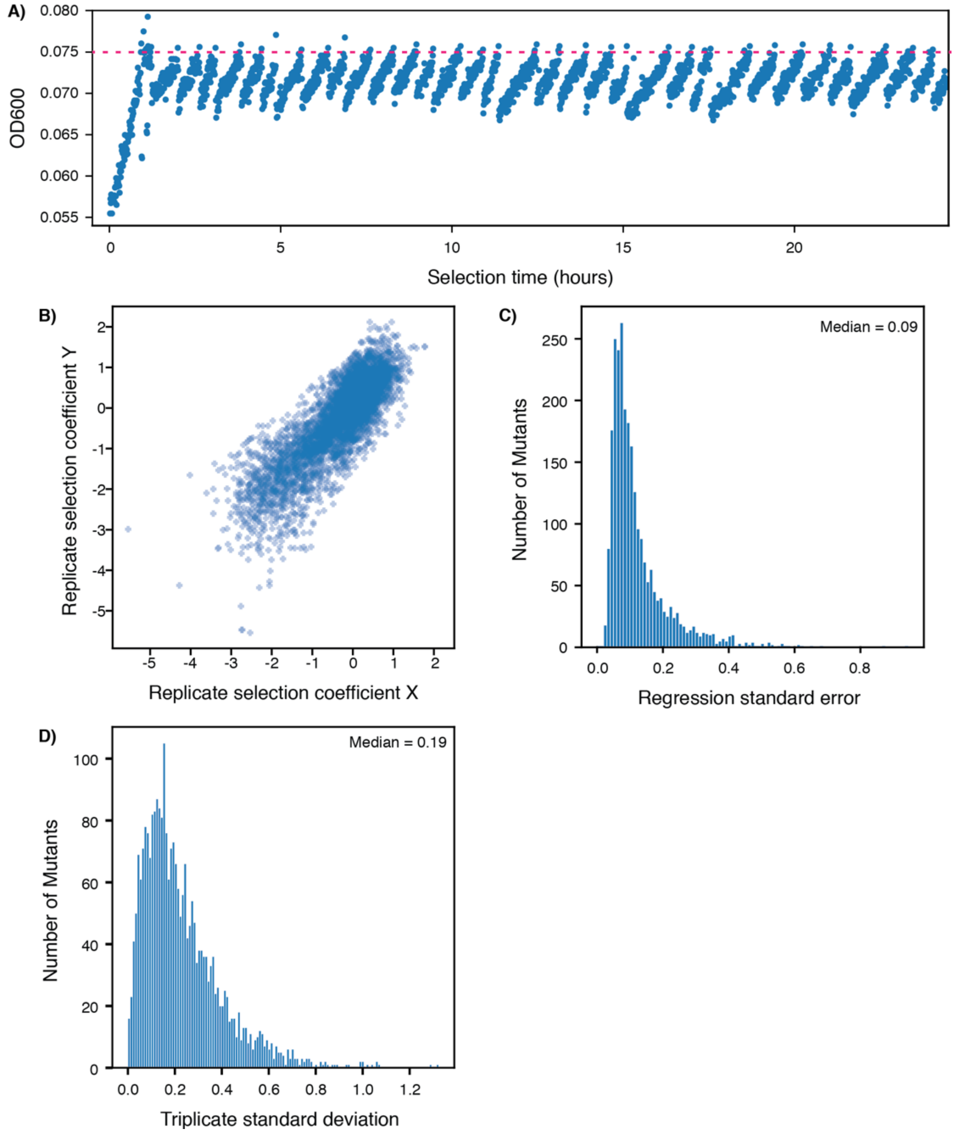


**Fig. S2. Determination of selection coefficients for DHFR.**
**A)** Example turbidostat trace from a selection experiment on a library of DHFR single point mutants. The OD600 value inferred from the voltage across an IR emitter-receiver pair is plotted as a function of time. The “clamp” OD value (0.075) is shown as a dashed red line. Decreases in OD correspond to dilution from automatic addition of M9 medium.
**B)** Comparison of all pairwise replicates for selection coefficients from triplicate deep mutational scanning on DHFR. The Pearson correlation $R^2$ value from linear regression was 0.70.
**C)** Distribution of standard errors for individual selection coefficients from a single replicate. Selection coefficients are the slope from a linear regression of allele frequency as a function of time in selection. The standard error here is the mean square of residuals.
**D)** Distribution of standard deviations for individual point mutants over replicate selection coefficients. Each mutant had a measured selection coefficient in at least 2 of the 3 replicates. The median of this distribution of standard deviations was used to determine the cut-offs for advantageous and disadvantageous mutations in **Figure 1**.

**Figure S3.**

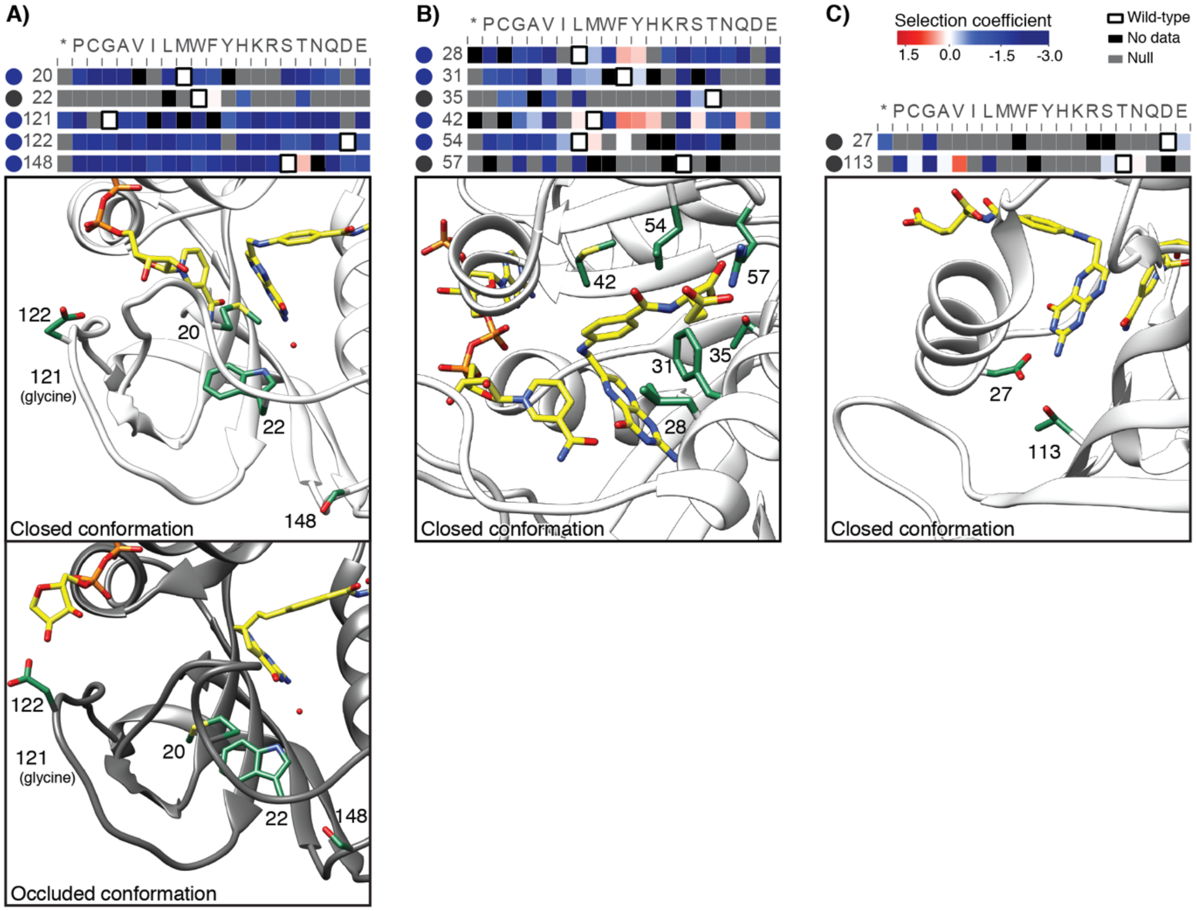


**Fig. S3. A-C) Residues previously known to have a functional role shown on the DHFR structure.** Important residues are colored green, labeled, and shown with slices of the –Lon heatmap (heatmap coloring by selection coefficient is as in **Figure 2**). The wild-type residue is outlined in black. A circle to the left of the heatmap indicates the mutational response category (colored as in **Figure 2C**). In A) the closed (upper, white, PDB ID: 3QL3) and occluded (lower, grey, PDB ID: 1RX4) conformation are shown to illustrate alternate stabilization of the two conformations by D122 (closed)(Miller & Benkovic, 1998) and S148 (occluded)(Miller, Wahnon, & Benkovic, 2001). For all other panels, only the closed conformation is shown.

64 **Figure S4.**

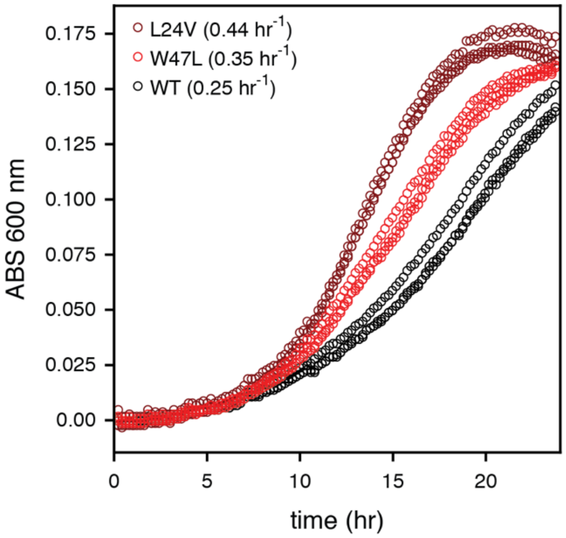


**Fig. S4. Growth curves for top advantageous mutations.** The absorbance (ABS) at 600 nm was monitored in 96-well plate format for monocultures of the selection strain transformed with strong advantageous mutants (L24V in dark red, W47L in bright red, wild-type in black). The doubling rates (top left in plot) were calculated from the early exponential phase of growth (see Methods). All growth curves are shown as sets of three biological replicates.

**Fig. S5.**

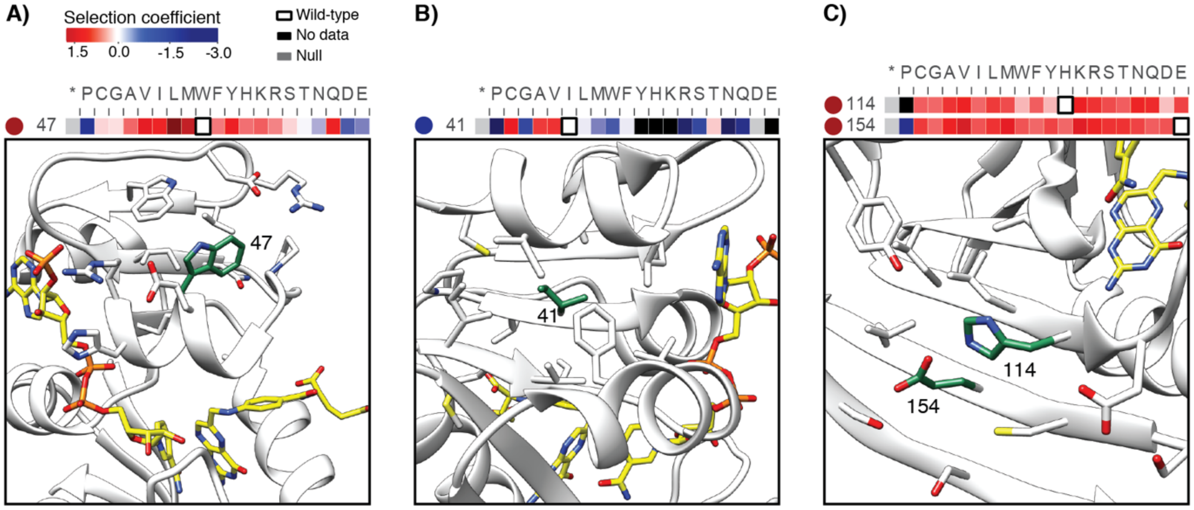


**Fig. S5. A-C) Example positions with multiple advantageous mutations hypothesized to be destabilizing, shown on the DHFR structure (PDB ID: 3QL3).** Wild-type residues are colored in green on the DHFR structure and depicted with slices of the –Lon heatmap (heatmap coloring is as in **Figure 2**). The wild-type residue is outlined in black on the heatmap. A circle to the left of the heatmap indicates the mutational response category (colored as in **Figure 2C**). In the examples here, advantageous mutations appear to disrupt core packing and a surface salt bridge.

**Figure S6.**

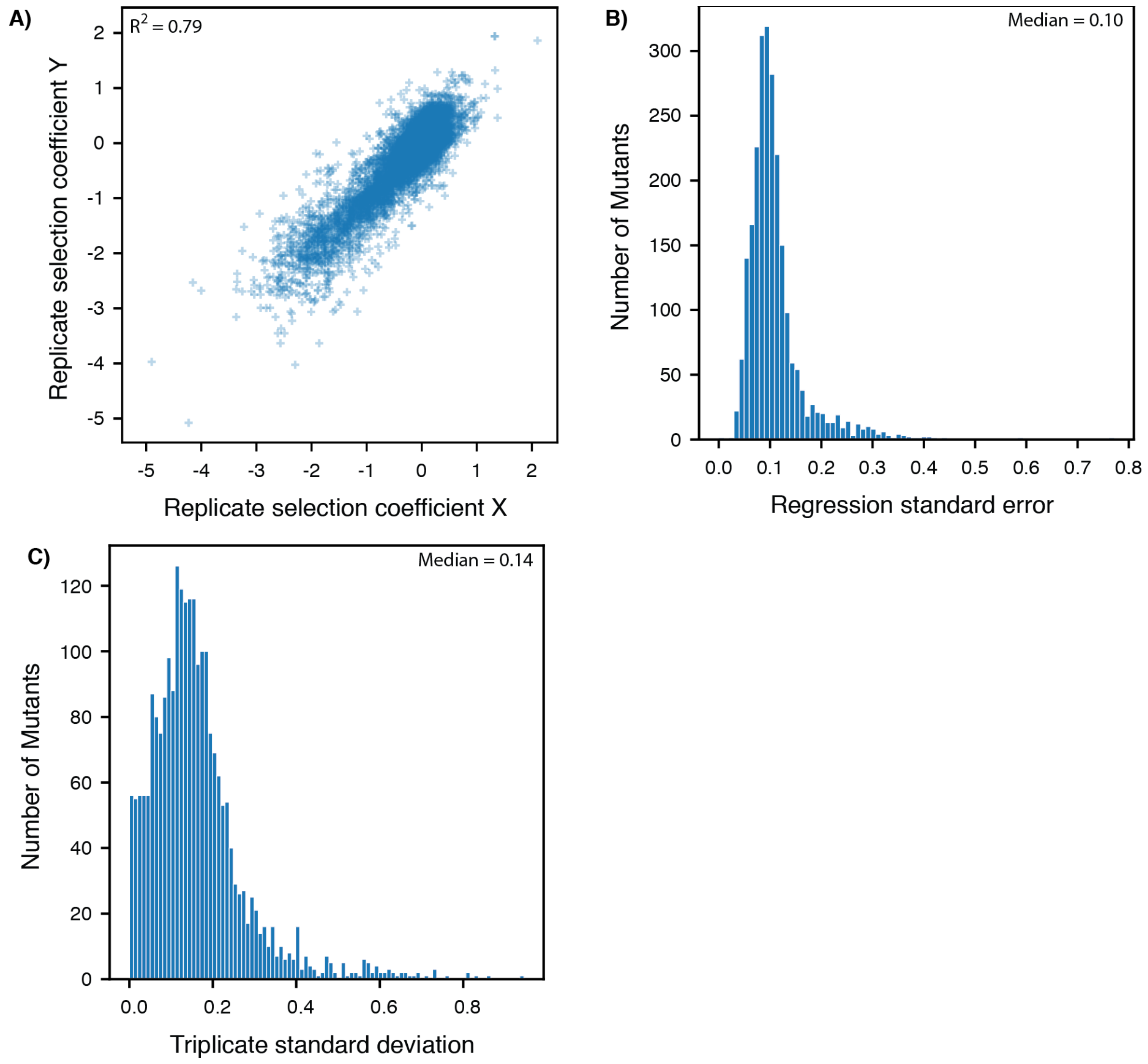


**Fig. S6. Quality of the selection under +Lon conditions. A)** Comparison of all pairwise replicates for +Lon selection coefficients from triplicate deep mutational scanning on DHFR. The Pearson correlation $R^2$ value from linear regression was 0.70.
**B)** Distribution of standard errors for individual +Lon selection coefficients from a single replicate. Selection coefficients are the slope from a linear regression of allele frequency as a function of time in selection. The standard error here is the mean square of residuals.
**C)** Distribution of standard deviations for individual point mutants over replicate +Lon selection coefficients. Each mutant had a measured selection coefficient in at least 2 of the 3 replicates.

**Figure S7.**

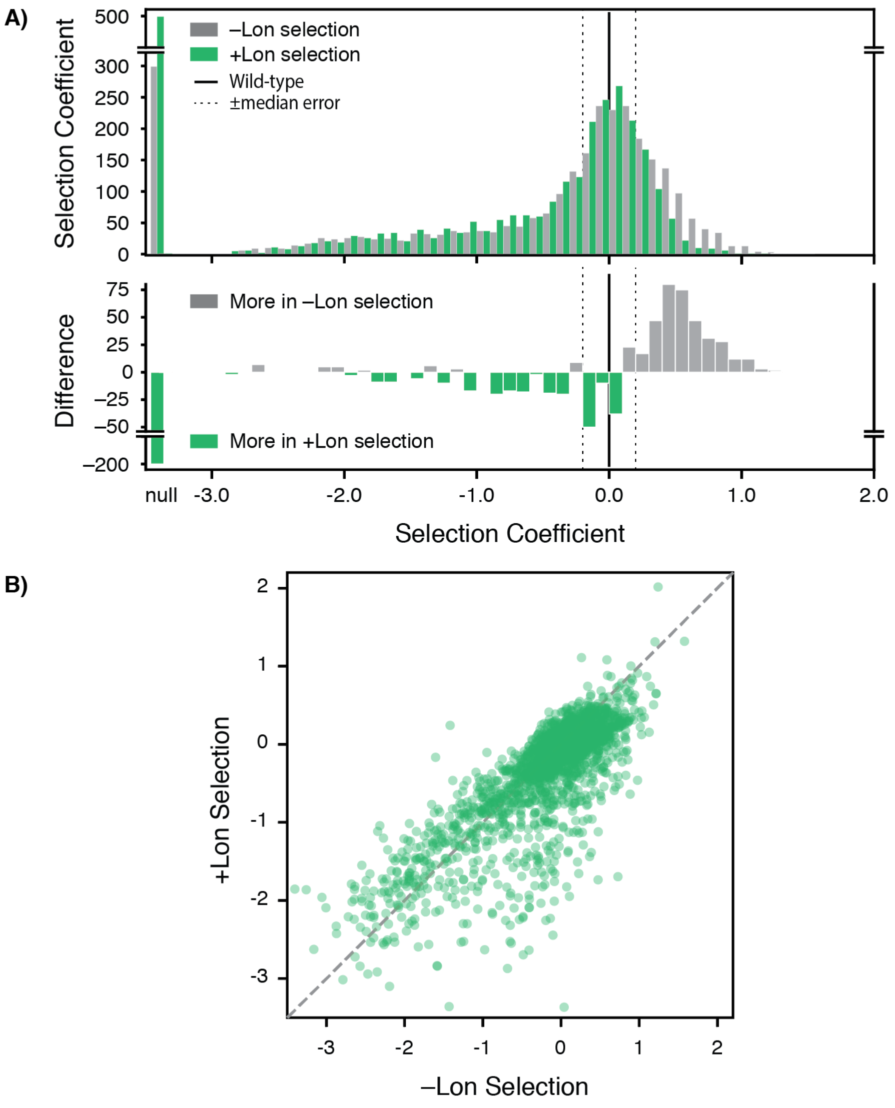


**Fig. S7. Comparison of selection coefficients ± Lon. A)** Histogram of selection coefficients (top) in –Lon (grey) and +Lon selection (green). The difference of the histograms (bottom) is shown with grey indicating more mutants for –Lon selection and green indicating more mutants for +Lon selection. The median of the standard deviation distribution (**Methods, Figure S2**) is indicated with dashed lines, as in **Figure 1**.
**B)** Scatterplot comparing selection coefficients in –Lon (grey) and +Lon selection (green) shows that mutations are generally repressed by Lon activity. Despite this general trend, we note that some top advantageous mutations are not impacted by Lon activity.

**Figure S8.**

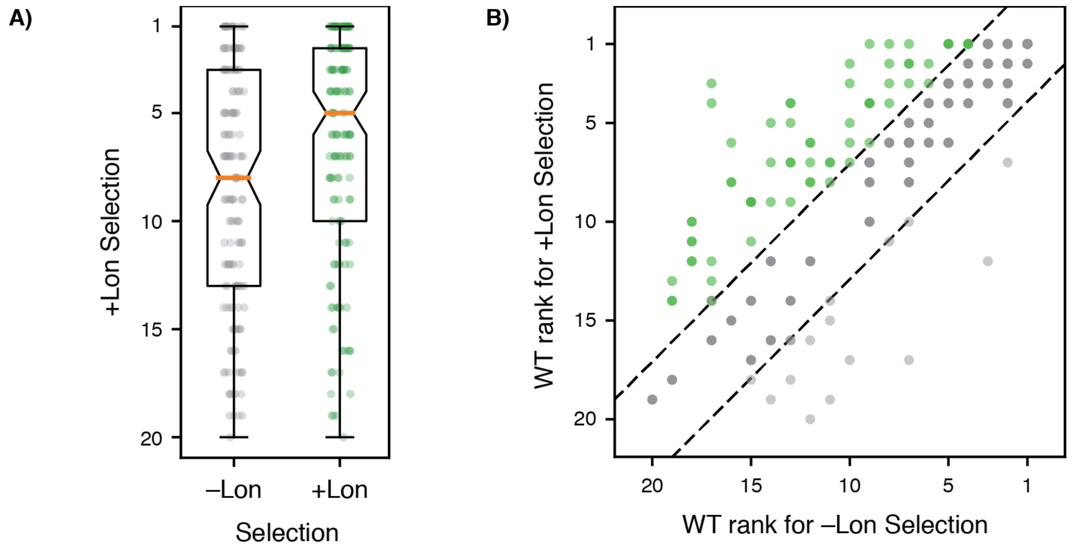


**Fig. S8. Ranks of the wild-type amino acid residues in ± Lon selections.**
**A)** Boxplot showing the distribution of wild-type amino acid residue rankings for –Lon (grey) and +Lon (green) selection. The wild-type amino acid residue ranking at each position is also shown as a distribution of points. Box plots show the median (orange bar) and upper/lower quartiles. The median wild-type amino acid residue rank is 8 for –Lon selection and 5 for +Lon selection.
**B)** Wild-type amino acid residue rankings from –Lon selection plotted against wild-type amino acid residue rankings from +Lon selection. Dashed lines show ±1 standard deviation for the change in rank between –Lon and +Lon selection.

**Figure S9.**

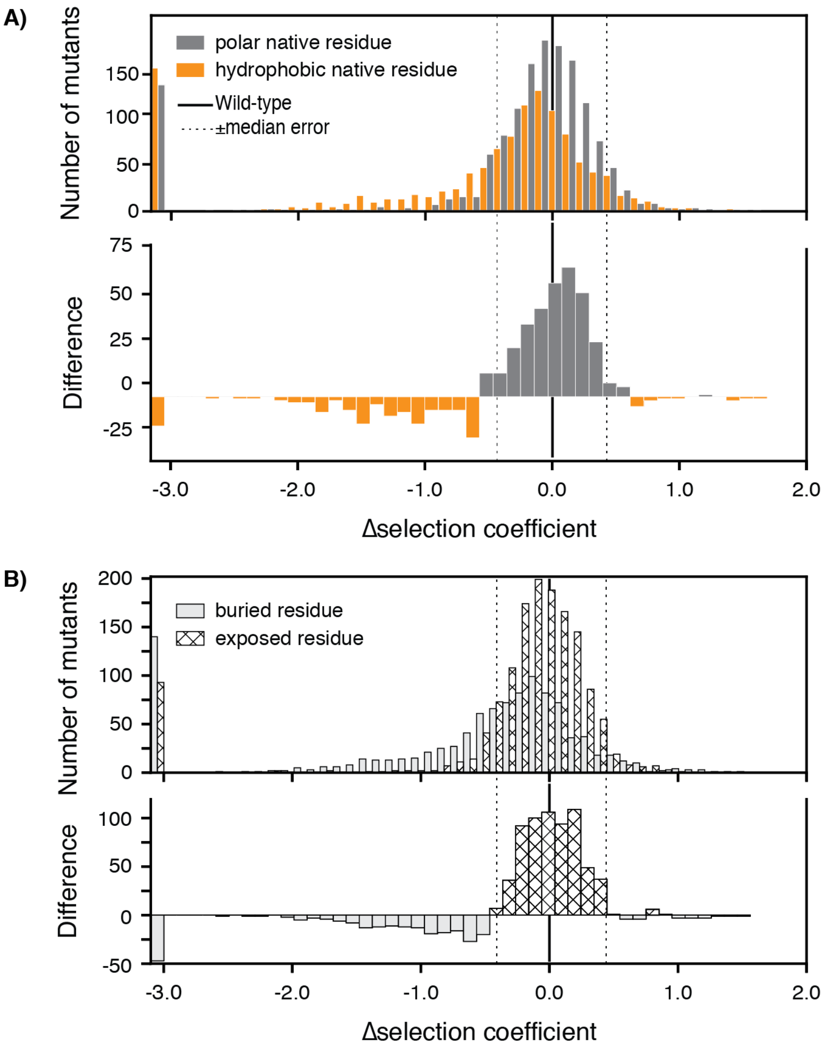


**Fig. S9. Δselection coefficients.**
**A)** Histogram of Δselection coefficients (top) with mutants at positions with hydrophobic (AVILMWFY) wild-type amino acid residues in orange and at positions with polar (HKRSTNQDE) wild-type amino acid residues in grey. Selection coefficients for positions with a wild-type P, G, or C residue are not included. The difference of the histograms (bottom) is shown with grey indicating more mutants to positions with a wild-type polar residue and orange indicating more mutants to positions with a wild-type hydrophobic residue. Dotted lines indicate twice the median of standard deviations from **Figure 1**.
**B)** Histogram of Δselection coefficients (top) with mutants at buried positions (solid) and at exposed positions (hatched) as listed in **Table S2**. Selection coefficients for positions that were Intolerant in –Lon selection are not included. The difference of the histograms (bottom) is shown with solid indicating more mutants to buried positions and hatched indicating more mutants to exposed positions.

**Figure S10.**

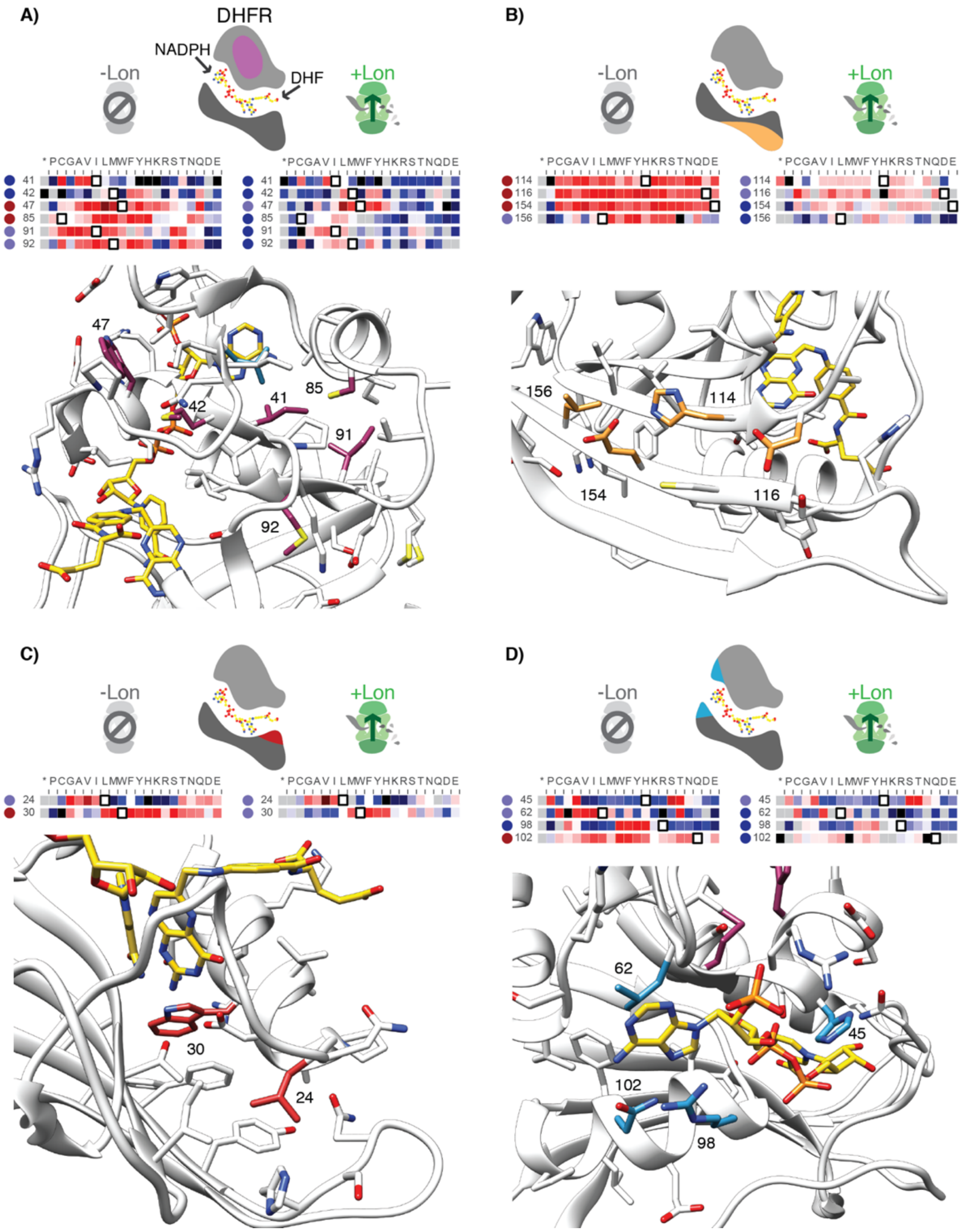


**Fig. S10. A-D) Structural context for hotspot residues from Figure 4**. For each panel, the hot spot region is indicated on a cartoon of DHFR: globular core in purple (**A**), the beta-sheet surface below the active site in gold (**B**), the base of the M20 loop in red (**C**) and the adenosine binding site in blue (**D**). Slices of the –Lon and +Lon heatmaps are shown for each position within the hot spot region (heatmap coloring is as in **Figure 2**). The wild-type residue is outlined in black. A circle to the left of the heatmap indicates the mutational response category (colored as in **Figure 2C,D**). For A-C) the structure shown is PDBID: 3QL3, and for D) the structure shown is PDB ID: 1RX1. In IRX1 (as in 1RX4), R98 is in proximity to the adenine ring. In 3QL3, R98 extends into bulk solvent. Residues within the hot spot cluster are labeled with their residue number.

**Figure S11.**

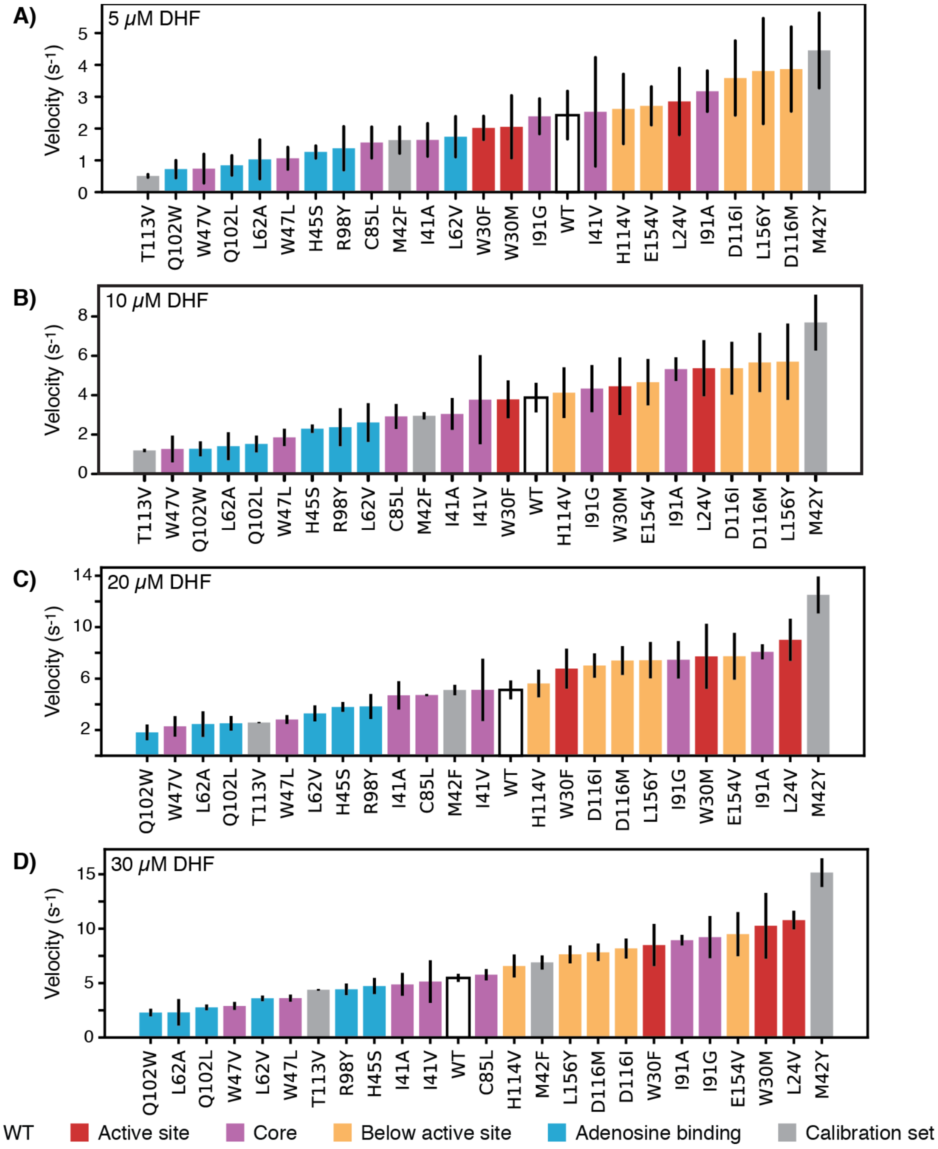


**Fig. S11.** ***In vitro*** **velocities of purified DHFR wild-type and point mutants.** Velocities were measured at A) 5, B) 10, C) 20, and D) 30 µM DHF. For each mutant, the bar is colored by the mutation's location within the hot spots from **Figure 4 and Figure S10**. Error bars represent ± 1 standard deviation from three independent experiments.

**Figure S12.**

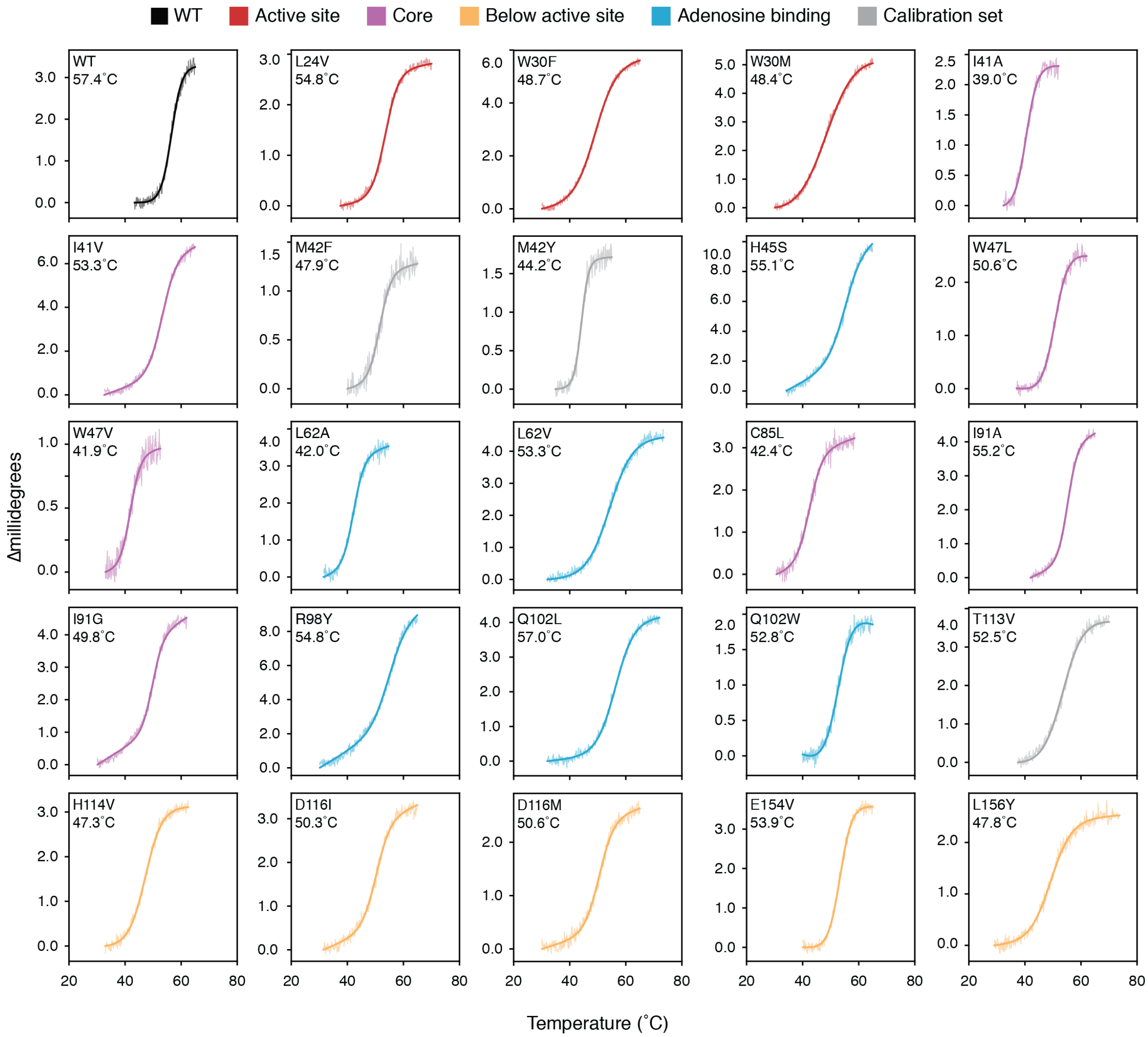


**Fig. S12. Thermal denaturation curves monitored by CD signal at 225m for selected hotspot mutants.** The curves are colored by hot spot cluster as in **Figure 4 and Figure S10**. The raw data are shown with thin lines and the fitted curves are shown as thick lines. For each plot, the mutant identity and apparent $T_M$ value are listed in the top left corner.

**Figure S13.**

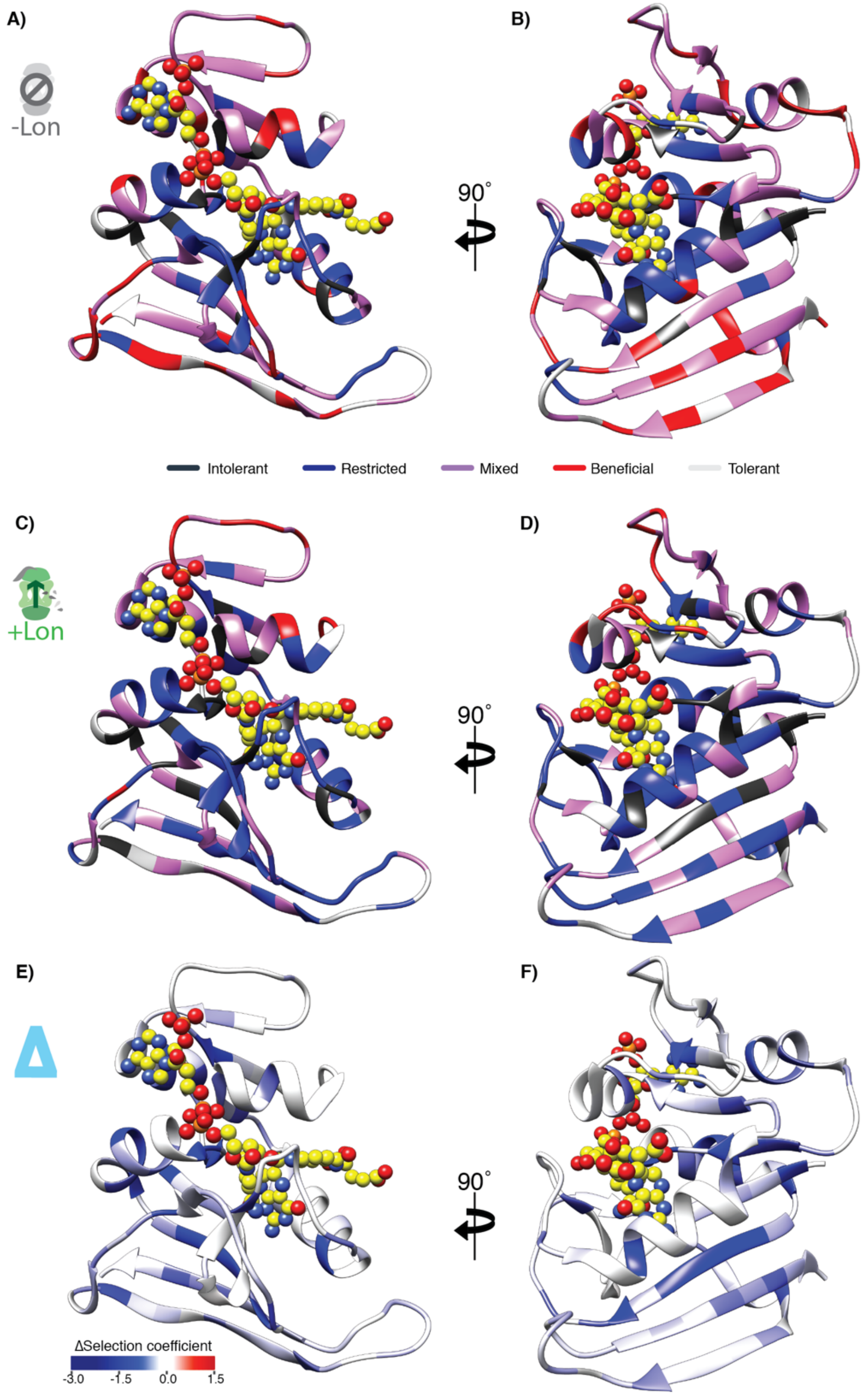


**Fig. S13. Selection coefficients under the two Lon expression regimes mapped on the DHFR structure.** Structural model of DHFR (PDB ID: 3QL3) in ribbon representation with the DHF substrate and the NADPH cofactor represented by spheres (yellow carbon and heteroatom coloring). (**A,B**) The residues are colored by mutational response category from **Figure 2C,D** for –Lon selection, (**C,D**) by mutational response category from +Lon selection, (**E,F**) or by the per-position mean Δselection coefficient from **Figure 3**.

**Figure S14.**

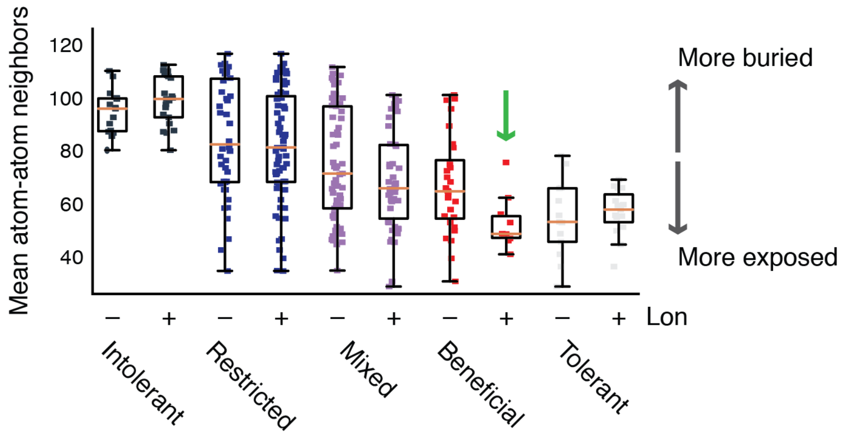


**Fig. S14. Burial of residues within each mutation response category reported as the mean number of atomic neighbors.** Each point represents one amino acid side chain, and the y-axis reports the average number of heavy atom neighbors within an 8 Å shell for all heavy atoms in that side chain. Box plots are overlaid on the distribution to show the median (orange bar) and upper/lower quartiles. Mutational response categories are shown for both –Lon and +Lon selection. The green arrow highlights the absence of buried Beneficial positions in +Lon selection.

**Figure S15.**

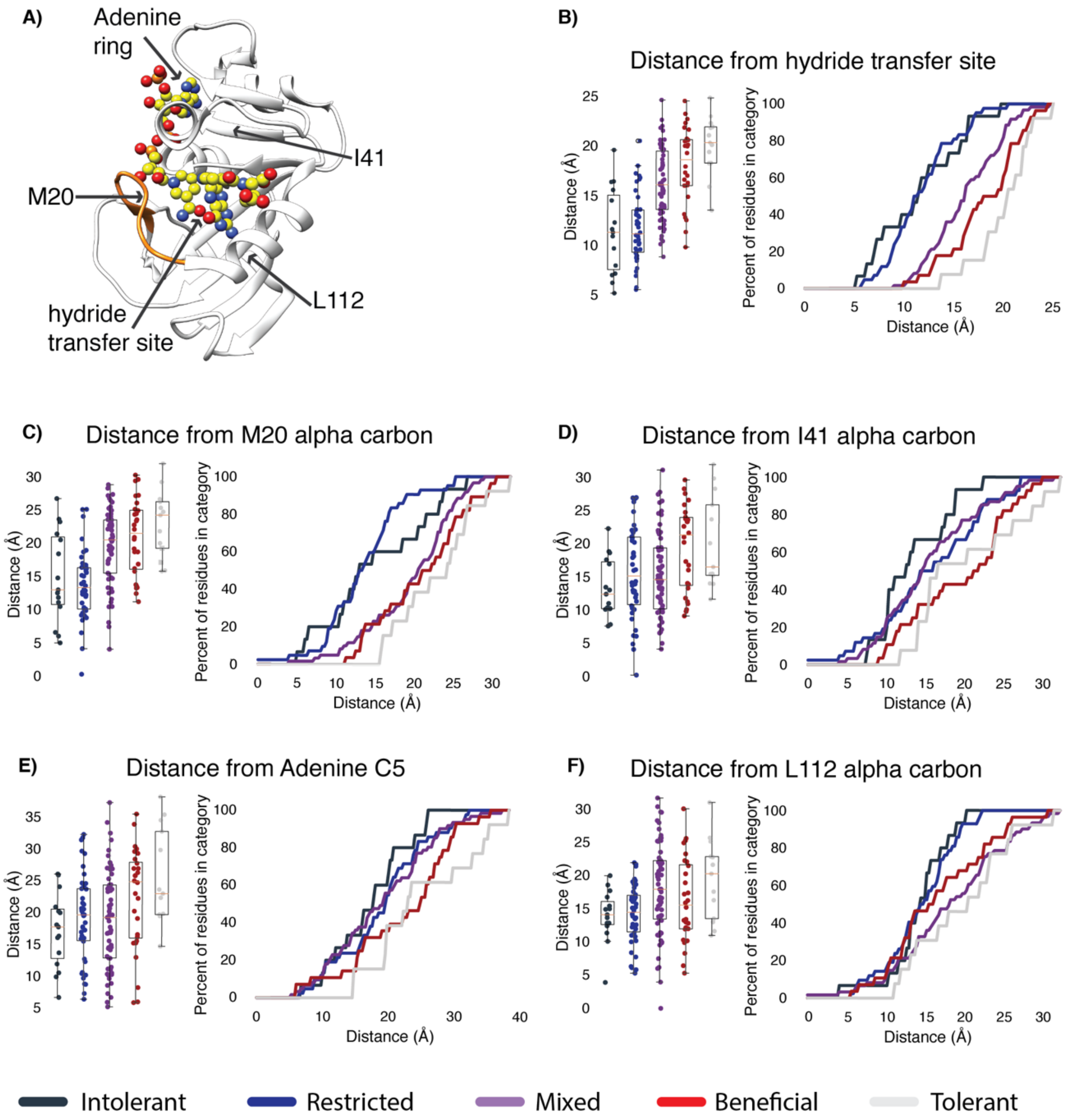


**Fig. S15. Residues in mutational response categories in the –Lon selection as a function of distance from several sites in the DHFR structure.**

**A)** Location of hydride transfer site, the M20 residue on the M20 loop (orange), and hot spot sites from **Figure 4** (the core of the globular domain represented by I41, the beta-sheet surface below the active site represented by L112, and the adenine ring on NADPH) indicated on the DHFR structure (PDB ID: 3QL3).

**B-F)** The distance relationships between each site and the residues in each mutational response category in the –Lon selection are shown (left) as boxplots with points representing the individual mutants and (right) as curves showing the percent of sequence positions in each mutational response category as a function of distance from the site. Boxplots and curves are colored by mutational response categories from –Lon selection as in **Figure 2C**.

**Table S1.**

Michaelis-Menten kinetics for the set of DHFR mutants used to calibrate the selection are reported together with the reference from which the values were taken.

| Mutant | $k_{cat}$ ($s^{-1}$) | $K_m$ ($\mu M$) | Reference |
|---|---|---|---|
| wt | 7.95 | 1.1 | Reynolds et al. Cell 2011(Reynolds, McLaughlin, & Ranganathan, 2011) |
| W22H | 1.89 | 18 | Reynolds et al. Cell 2011 |
| L28F | 18.5 | 9.9 | This work |
| L28Y | 19.2 | 21.2 | This work |
| F31Y | 20.61 | 80 | Reynolds et al. Cell 2011 |
| F31V | 8.65 | 108 | Reynolds et al. Cell 2011 |
| M42F | 79.2 | 13 | Reynolds et al. Cell 2011 |
| L54F | 6.3 | 0.7 | Huang et al. Biochemistry 1994(Huang, Wagner, & Benkovic, 1994) |
| L54I | 7.88 | 35 | Reynolds et al. Cell 2011 |
| T113V | 32.9 | 21.4 | Fierke and Benkovic Biochemistry 1989(Fierke & Benkovic, 1989) |
| G121V | 0.3 | 6.1 | Reynolds et al. Cell 2011 |
| F31Y/L54I | 1.94 | 168.3 | Reynolds et al. Cell 2011 |
| F31Y/G121V | 0.13 | 90.6 | Reynolds et al. Cell 2011 |
| M42F/G121V | 0.4 | 71.8 | Reynolds et al. Cell 2011 |
| L54I/G121V | 0.22 | 73 | Reynolds et al. Cell 2011 |

**Table S2.**

Burial classification for DHFR positions from the Getarea server(Fraczkiewicz & Braun, 1998) as described in **Methods**.

| Pos. | –Lon | +Lon | >20% SASA |
|---|---|---|---|
| 1 | Intol. | Intol. | Bur. |
| 2 | Intol. | Intol. | Bur. |
| 3 | Dele. | Dele. | Bur. |
| 4 | Dele. | Dele. | Bur. |
| 5 | Dele. | Dele. | Bur. |
| 6 | Dele. | Dele. | Bur. |
| 7 | Dele. | Dele. | Bur. |
| 8 | Dele. | Dele. | Bur. |
| 9 | Dele. | Dele. | Bur. |
| 10 | Dele. | Dele. | Exp. |
| 11 | Mixed | Mixed | Exp. |
| 12 | Mixed | Mixed | Exp. |
| 13 | Dele. | Dele. | Bur. |
| 14 | Intol. | Dele. | Bur. |
| 15 | Intol. | Intol. | Bur. |
| 16 | Dele. | Dele. | Exp. |
| 17 | Dele. | Dele. | Exp. |
| 18 | Mixed | Dele. | Exp. |
| 19 | Mixed | Mixed | Exp. |
| 20 | Dele. | Dele. | Bur. |
| 21 | Dele. | Dele. | Bur. |
| 22 | Intol. | Intol. | Bur. |
| 23 | Mixed | Dele. | Exp. |
| 24 | Mixed | Mixed | Bur. |
| 25 | Mixed | Tol. | Exp. |
| 26 | Dele. | Dele. | Exp. |
| 27 | Intol. | Intol. | Bur. |
| 28 | Dele. | Dele. | Exp. |
| 29 | Dele. | Dele. | Exp. |
| 30 | Bene. | Mixed | Bur. |
| 31 | Dele. | Dele. | Exp. |
| 32 | Dele. | Dele. | Exp. |
| 33 | Mixed | Mixed | Exp. |
| 34 | Mixed | Dele. | Bur. |
| 35 | Intol. | Intol. | Bur. |
| 36 | Mixed | Mixed | Exp. |
| 37 | Mixed | Mixed | Exp. |
| 38 | Mixed | Dele. | Exp. |
| 39 | Mixed | Mixed | Bur. |
| 40 | Mixed | Dele. | Bur. |
| 41 | Dele. | Dele. | Bur. |
| 42 | Dele. | Dele. | Bur. |
| 43 | Intol. | Intol. | Bur. |
| 44 | Mixed | Mixed | Bur. |
| 45 | Mixed | Mixed | Exp. |
| 46 | Intol. | Intol. | Bur. |
| 47 | Bene. | Mixed | Bur. |
| 48 | Bene. | Bene. | Exp. |
| 49 | Dele. | Dele. | Bur. |
| 50 | Dele. | Dele. | Bur. |
| 51 | Mixed | Tol. | Exp. |
| 52 | Tol. | Bene. | Exp. |
| 53 | Mixed | Bene. | Exp. |
| 54 | Dele. | Dele. | Bur. |
| 55 | Tol. | Bene. | Exp. |
| 56 | Mixed | Tol. | Exp. |
| 57 | Intol. | Dele. | Bur. |
| 58 | Mixed | Tol. | Exp. |
| 59 | Mixed | Mixed | Bur. |
| 60 | Mixed | Dele. | Bur. |
| 61 | Dele. | Intol. | Bur. |
| 62 | Mixed | Dele. | Bur. |
| 63 | Mixed | Dele. | Bur. |
| 64 | Mixed | Bene. | Exp. |
| 65 | Mixed | Bene. | Exp. |
| 66 | Mixed | Mixed | Exp. |
| 67 | Mixed | Bene. | Exp. |
| 68 | Bene. | Bene. | Exp. |
| 69 | Mixed | Mixed | Bur. |
| 70 | Tol. | Bene. | Exp. |
| 71 | Mixed | Mixed | Exp. |
| 72 | Bene. | Mixed | Bur. |
| 73 | Mixed | Mixed | Exp. |
| 74 | Mixed | Dele. | Bur. |
| 75 | Mixed | Mixed | Bur. |
| 76 | Mixed | Mixed | Exp. |
| 77 | Bene. | Mixed | Exp. |
| 78 | Bene. | Mixed | Bur. |
| 79 | Mixed | Mixed | Exp. |
| 80 | Mixed | Mixed | Exp. |
| 81 | Dele. | Intol. | Bur. |
| 82 | Mixed | Dele. | Exp. |
| 83 | Tol. | Mixed | Exp. |
| 84 | Mixed | Mixed | Bur. |
| 85 | Bene. | Dele. | Bur. |
| 86 | Bene. | Tol. | Exp. |
| 87 | Tol. | Mixed | Exp. |
| 88 | Bene. | Tol. | Exp. |
| 89 | Mixed | Tol. | Exp. |
| 90 | Dele. | Dele. | Bur. |
| 91 | Mixed | Dele. | Bur. |
| 92 | Mixed | Dele. | Bur. |
| 93 | Dele. | Intol. | Bur. |
| 94 | Dele. | Intol. | Bur. |
| 95 | Intol. | Intol. | Bur. |
| 96 | Intol. | Intol. | Exp. |
| 97 | Dele. | Dele. | Bur. |
| 98 | Dele. | Dele. | Exp. |
| 99 | Dele. | Dele. | Bur. |
| 100 | Dele. | Dele. | Bur. |
| 101 | Mixed | Mixed | Exp. |
| 102 | Bene. | Dele. | Exp. |
| 103 | Mixed | Dele. | Bur. |
| 104 | Intol. | Intol. | Bur. |
| 105 | Tol. | Tol. | Exp. |
| 106 | Mixed | Mixed | Bur. |
| 107 | Intol. | Intol. | Bur. |
| 108 | Tol. | Mixed | Exp. |
| 109 | Mixed | Mixed | Bur. |
| 110 | Dele. | Intol. | Bur. |
| 111 | Mixed | Dele. | Bur. |
| 112 | Mixed | Intol. | Bur. |
| 113 | Intol. | Intol. | Bur. |
| 114 | Bene. | Mixed | Exp. |
| 115 | Mixed | Dele. | Bur. |
| 116 | Bene. | Mixed | Bur. |
| 117 | Dele. | Dele. | Bur. |
| 118 | Bene. | Dele. | Exp. |
| 119 | Mixed | Mixed | Bur. |
| 120 | Bene. | Mixed | Exp. |
| 121 | Dele. | Dele. | Bur. |
| 122 | Dele. | Dele. | Exp. |
| 123 | Dele. | Dele. | Exp. |
| 124 | Mixed | Mixed | Exp. |
| 125 | Dele. | Intol. | Bur. |
| 126 | Bene. | Dele. | Bur. |
| 127 | Bene. | Bene. | Exp. |
| 128 | Mixed | Dele. | Bur. |
| 129 | Bene. | Mixed | Exp. |
| 130 | Bene. | Tol. | Exp. |
| 131 | Mixed | Dele. | Exp. |
| 132 | Bene. | Mixed | Exp. |
| 133 | Dele. | Intol. | Bur. |
| 134 | Bene. | Tol. | Exp. |
| 135 | Bene. | Mixed | Exp. |
| 136 | Tol. | Tol. | Exp. |
| 137 | Bene. | Mixed | Exp. |
| 138 | Mixed | Mixed | Exp. |
| 139 | Tol. | Dele. | Exp. |
| 140 | Bene. | Mixed | Exp. |
| 141 | Mixed | Dele. | Exp. |
| 142 | Bene. | Tol. | Exp. |
| 143 | Tol. | Tol. | Exp. |
| 144 | Mixed | Dele. | Exp. |
| 145 | Tol. | Tol. | Exp. |
| 146 | Tol. | Mixed | Exp. |
| 147 | Dele. | Dele. | Bur. |
| 148 | Dele. | Dele. | Exp. |
| 149 | Dele. | Dele. | Exp. |
| 150 | Mixed | Dele. | Exp. |
| 151 | Mixed | Dele. | Bur. |
| 152 | Bene. | Mixed | Bur. |
| 153 | Mixed | Dele. | Bur. |
| 154 | Bene. | Mixed | Bur. |
| 155 | Mixed | Dele. | Bur. |
| 156 | Mixed | Dele. | Bur. |
| 157 | Mixed | Mixed | Exp. |
| 158 | Tol. | Dele. | Exp. |
| 159 | Bene. | Tol. | Exp. |

**Table S3.**

*In vitro* turnover for selected advantageous measured as described in **Methods** at multiple concentrations of DHF are reported with the standard deviation over three independent experiments.

| | 5 μM DHF | | 10 μM DHF | | 20 μM DHF | | 30 μM DHF | |
|---|---|---|---|---|---|---|---|---|
| Mutant | Velocity | St. dev. | Velocity | St. dev. | Velocity | St. dev. | Velocity | St. dev. |
| _wt | 2.42 | 0.75 | 3.88 | 0.75 | 5.12 | 0.74 | 5.48 | 0.37 |
| L24V | 2.85 | 1.05 | 5.37 | 1.43 | 9.01 | 1.64 | 10.79 | 0.86 |
| W30F | 2.02 | 0.37 | 3.79 | 0.96 | 6.77 | 1.56 | 8.50 | 1.94 |
| W30M | 2.05 | 0.98 | 4.45 | 1.46 | 7.73 | 2.53 | 10.27 | 3.03 |
| I41A | 1.64 | 0.52 | 3.05 | 0.81 | 4.69 | 1.11 | 4.88 | 1.06 |
| I41V | 2.52 | 1.71 | 3.77 | 2.27 | 5.11 | 2.44 | 5.13 | 1.96 |
| M42F | 3.08 | 1.40 | 4.62 | 1.68 | 6.42 | 1.24 | 6.90 | 0.64 |
| M42Y | 4.45 | 1.18 | 7.69 | 1.41 | 12.51 | 1.43 | 15.16 | 1.32 |
| H45S | 1.26 | 0.19 | 2.29 | 0.21 | 3.79 | 0.38 | 4.73 | 0.74 |
| W47L | 1.24 | 0.35 | 2.12 | 0.36 | 2.91 | 0.39 | 4.02 | 0.00 |
| W47V | 0.74 | 0.45 | 1.27 | 0.68 | 2.28 | 0.79 | 2.90 | 0.37 |
| L62A | 1.03 | 0.61 | 1.41 | 0.71 | 2.46 | 1.00 | 2.30 | 1.22 |
| L62V | 1.99 | 0.54 | 3.06 | 0.68 | 3.66 | 0.41 | 3.85 | 0.00 |
| C85L | 1.56 | 0.49 | 2.92 | 0.63 | 4.71 | 0.09 | 5.77 | 0.52 |
| I91A | 3.17 | 0.64 | 5.32 | 0.60 | 8.08 | 0.59 | 8.94 | 0.49 |
| I91G | 2.38 | 0.55 | 4.33 | 1.20 | 7.46 | 1.46 | 9.22 | 1.95 |
| R98Y | 1.38 | 0.68 | 2.38 | 0.97 | 3.83 | 0.97 | 4.43 | 0.53 |
| Q102L | 0.84 | 0.31 | 1.52 | 0.43 | 2.52 | 0.57 | 2.77 | 0.26 |
| Q102W | 0.72 | 0.28 | 1.27 | 0.38 | 1.81 | 0.61 | 2.29 | 0.35 |
| T113V | 0.51 | 0.05 | 1.19 | 0.09 | 2.58 | 0.04 | 4.38 | 0.08 |
| H114V | 2.50 | 1.24 | 3.91 | 1.42 | 5.26 | 1.01 | 5.90 | 0.58 |
| D116I | 3.31 | 1.23 | 4.92 | 1.26 | 6.58 | 0.68 | 7.55 | 0.22 |
| D116M | 3.63 | 1.46 | 5.35 | 1.62 | 6.95 | 0.93 | 7.28 | 0.25 |
| E154V | 2.71 | 0.60 | 4.66 | 1.18 | 7.73 | 1.82 | 9.50 | 2.02 |
| L156Y | 3.80 | 1.66 | 5.70 | 1.94 | 7.43 | 1.41 | 7.64 | 0.82 |

**Table S4.**

Apparent $T_m$ values from thermal denaturation experiments monitored by CD signal at 225 nm are reported along with the Δselection coefficient (Lon impact) value depicted in **Figure 4D**.

| Mutant | $T_m$ [deg C] | Δselection coefficient |
|---|---|---|
| _wt | 57.4 | 0 |
| L24V | 54.8 | 0.72 |
| W30F | 48.7 | -0.15 |
| W30M | 48.4 | -0.37 |
| I41A | 39.0 | -1.24 |
| I41V | 53.3 | -0.07 |
| M42F | 47.9 | -0.29 |
| M42Y | 44.2 | -0.1 |
| H45S | 55.1 | -0.33 |
| W47L | 50.6 | -0.2 |
| W47V | 41.9 | -1.46 |
| L62A | 42.0 | -1.35 |
| L62V | 53.3 | -0.21 |
| C85L | 42.4 | -0.78 |
| I91A | 55.2 | -0.6 |
| I91G | 49.8 | -0.9 |
| R98Y | 54.8 | -0.47 |
| Q102L | 57.0 | -0.3 |
| Q102W | 52.8 | -0.69 |
| T113V | 52.5 | -0.85 |
| H114V | 47.3 | -0.66 |
| D116I | 50.3 | -0.34 |
| D116M | 50.6 | -0.72 |
| E154V | 53.9 | -0.9 |
| L156Y | 47.8 | -0.65 |

**Source data 1. (separate file)**

Selection coefficients for –Lon selection measured as described in **Methods** are reported with the standard deviation between biological replicates and the standard error from linear regression (as calculated by Enrich2(Rubin et al., 2017)). Values are reported as calculated, but based on the selection calibration, differences between selection coefficients with values below ~ –2.5 are not interpretable.

**Source data 2. (separate file)**

Selection coefficients for +Lon selection measured as described in **Methods** are reported with the standard deviation between biological replicates and the standard error from linear regression. Values are reported as calculated, but based on the selection calibration, differences between selection coefficients with values below ~ –2.5 are not interpretable.

**Source data 3. (separate file)**

Plasmid descriptions and primer sequences for all constructs used in the experiments.

**Source data 4. (separate file)**

Raw deep sequencing counts for the calibration set of mutants –Lon selection. Counts are recorded for all turbidostat timepoints over 3 repeats.

**Source data 5. (separate file)**

Raw deep sequencing counts for single point mutants in –Lon selection. Counts are recorded for a sample collected from the transformation rescue medium, a sample from overnight outgrowth in supplemented M9, and for all turbidostat timepoints over 6 experiments. In each experiment, 2 of 4 sublibraries were screened as described in **Methods**, for a total of 3 repeats over the full library.

**Source data 6. (separate file)**

Raw deep sequencing counts for single point mutants in +Lon selection. Counts are recorded as in **Source data 5**.
